# Paracrine signaling by pancreatic δ cells determines the glycemic set point in mice

**DOI:** 10.1101/2022.06.29.496132

**Authors:** Jessica L Huang, Sharon Lee, Mohammad S Pourhosseinzadeh, Niels Krämer, Jaresley V Guillen, Naomi H Cinque, Paola A Aniceto, Shinichiro Koike, Mark O Huising

## Abstract

Pancreatic islets contain several endocrine cell types that coordinate to maintain blood glucose homeostasis. While β and α cells are thought to be the main drivers of glucose homeostasis through insulin and glucagon secretion respectively, the contribution of δ cells and somatostatin (SST) secretion to establishing the glycemic set point remains unresolved. Here we remove local SST signaling from δ cells within the pancreatic islet to investigate their contribution to the glycemic set point. Our data demonstrate that ablating δ cells or SST leads to a sustained decrease in the glycemic set point. This coincides with a decreased glucose threshold for insulin response from β cells, leading to increased insulin secretion to the same glucose challenge. In contrast, α cell ablation had no effect on glycemic set point. Collectively, these data establish the physiological role of δ cells in determining the glycemic set point through their interaction with β cells.

## INTRODUCTION

Blood glucose levels are maintained within a narrow range around the glycemic set point, defined as a fixed level of blood glucose that the body aims to achieve in between meals (Matschinsky & Davis 1998). Glucose homeostasis changes during postnatal development in rodents, but is generally stable within an adult individual throughout the lifespan unless disrupted by disease. In humans, this set point is around 90 mg/dL (approximately 5 mM) (Gerich 1993), while in mice it is around 120-140 mg/dL (approximately 7-8 mM) (Ewing & Tauber 1964; Blum *et al*. 2012; Rodriguez-Diaz *et al*. 2018). Tight regulation of blood glucose homeostasis is crucial, as chronic hyperglycemia causes a plethora of long-term complications, while hypoglycemia is acutely lifethreatening.

The hormones released by pancreatic islets are known to play critical roles in blood glucose homeostasis. Under prandial conditions when glycemia is high, β cells within the islet secrete insulin to signal for the uptake and storage of glucose. Conversely, under post-prandial conditions when glycemia is low, α cells secrete glucagon to stimulate hepatic glucose production. There is increasing evidence that paracrine glucagon signaling also amplifies glucose-stimulated insulin secretion (GSIS) by direct stimulation of β cells (Chambers *et al*. 2017; Svendsen *et al*. 2018; Capozzi *et al*. 2019a, b; Zhu *et al*. 2019; Tellez *et al*. 2020; Liu *et al*. 2021). This suggests that glucagon acts locally to stimulate GSIS as a paracrine factor during the prandial state, while systemic glucagon action in the absence of insulin release is responsible for its counterregulatory function during the post-prandial state (Huising 2020). The third major cell type of the islet is the δ cell, which releases somatostatin (SST) to inhibit both β and α cells.

The glycemic set point is often attributed to the crossover point between β and α cell glucose response (Pagliara *et al*. 1974; Rodriguez-Diaz *et al*. 2018). However, the question of where and how the glycemic set point is determined continues to elicit debate. The central nervous system (CNS) is a key regulator of blood glucose homeostasis that has been proposed to be responsible for the glycemic set point (Alonge *et al*. 2021). Indeed, glucose-sensing neurons are present at various regions of the brain, including the ventromedial nucleus, paraventricular nucleus, and lateral hypothalamus (Thorens 2012). Moreover, defects in glucose sensing at these sites contributes to type 2 diabetes. It has also been demonstrated that the CNS is capable of lowering glycemia in an insulin-independent manner (German *et al*. 2011; Morton *et al*. 2013; Ryan *et al*. 2013).

While these observations clearly indicate that the CNS plays an important role in controlling glucose levels, there is limited evidence to indicate that CNS sites determine the plasma glucose levels in between meals. In contrast, there is compelling evidence that the pancreatic islet is the major glucostat of the body. Grafting islets from different donor species into diabetic nude mice causes recipients to re-establish a set point matching that of the donors, demonstrating that islets are both sufficient to restore normoglycemia and responsible for determining the homeostatic set point of glucose (Carroll *et al*. 1992). More recently, an experiment where human, mouse or non-human islets were grafted into the anterior chamber of the eye in diabetic nude mice demonstrated that the glycemic set point established in the recipient mice was the same as that of the donor species and was dependent on paracrine interactions within the grafted islets (Rodriguez-Diaz *et al*. 2018). These experiments provide evidence that paracrine signals within the pancreatic islet are key players in establishing the glycemic set point, but did not address the potential paracrine contribution of δ cells to this set point.

β cells in mice do not respond to glucose until around 7-8 mM glucose (Vieira *et al*. 2007), and α cells in mice are maximally suppressed at 7 mM glucose (Vieira *et al*. 2007; Lai *et al*. 2018). Therefore, at the glycemic set point, β cells are not yet activated, and α cell activity is at a nadir. In contrast, δ cells are active over a range of glucose levels and are known to inhibit β and α cells in a paracrine manner (Hauge-Evans *et al*. 2009; van der Meulen *et al*. 2015; Lai *et al*. 2018; Xu *et al*. 2020; Singh *et al*. 2021). While knockout of SST has been shown to augment glucose-stimulated insulin secretion and arginine-stimulated glucagon secretion, the physiological contribution of δ cell-secreted SST to the glycemic set point has not been established.

There is evidence that indirectly points to a role of δ cell-derived SST in determining the glycemic set point through their communication with β cells. Our lab has established that β cells co-secrete the hormone Urocortin 3 (UCN3) with insulin, and that within the islet UCN3 acts solely on δ cells to stimulate SST secretion in a paracrine manner (van der Meulen *et al*. 2015). Onset of UCN3 expression at around 2 weeks of age is associated with an increase in the glycemic set point in mice at around 2 weeks of age (Blum *et al*. 2012; van der Meulen *et al*. 2012). Premature induction of UCN3 in neonatal mice caused a comparable and premature increase in glycemia, while continued induction of UCN3 after endogenous UCN3 expression has been established had no further effect, demonstrating that the increased set point observed in young mice is caused by the onset of UCN3 expression. UCN3 expression is also known to be downregulated in Type 2 Diabetes (T2D), which leads to reduced SST secretion to allow for a compensatory increase in insulin in the face of peripheral insulin resistance (Blum *et al*. 2014; van der Meulen *et al*. 2015; Kavalakatt *et al*. 2019). Restoring UCN3 expression and the ensuing SST feedback that suppresses insulin secretion in diabetic *ob/ob* mice aggravated hyperglycemia (van der Meulen *et al*. 2015). From these observations, we predicted that δ cell feedback via the local release of endogenous SST helps determine the glycemic set point (Huising *et al*. 2018).

Here we set out to rigorously test the hypothesis that SST contributes to the glycemic set point. We do so with three complementary mouse models of removing SST-mediated inhibition of β cells in adult mice that consistently leads to an immediate and sustained decrease of 20-30 mg/dL in blood glucose. We demonstrate that the effect on the glycemic set point is specific to the loss of pancreatic δ cell-derived SST, ruling out contributions of non-pancreatic sources of SST to this phenotype. We then demonstrate that this acute drop in the glycemic set point is due to an increase in insulin secretion by measuring plasma insulin *in vivo* and secreted insulin *ex vivo*. Parallel pancreatic α cell ablation experiments do not shift the glycemic set point, confirming prior published observations (Thorel *et al*. 2011; Pedersen *et al*. 2013; Shiota *et al*. 2013). Furthermore, quantifying the calcium responses of several thousand β cells within intact islets over time revealed a decreased glucose threshold for β cell response of approximately 1 mM in the absence of δ cells. This reduction of the β cell glucose threshold closely matches the 20-30 mg/dL reduction in the glycemic set point observed *in vivo* upon interruption of δ cell function in multiple parallel experiments. We conclude that δ cells shift the glycemic set point through their local inhibitory interactions with β cells, modulating their glucose threshold for insulin secretion.

## RESULTS

### Absence of somatostatin lowers the glycemic set point

To investigate the contribution of SST to the glycemic set point, we first used mice with the *Sst*-IRES-Cre transgene, which is known to disrupt the expression of SST (Viollet *et al*. 2017). Using homozygous *Sst*-IRES-Cre mice (*Sst*-Cre^TG/TG^) crossed to a floxed YFP reporter (lsl-YFP) allowed for identification of δ cells with YFP and confirmed that SST is absent but δ cells remain in *Sst*-Cre^TG/TG^ mice. In contrast, SST remains present in δ cells of heterozygous littermates (*Sst*-Cre^+/TG^ x lsl-YFP) (Figure 1A and 1B). The absence of *Sst* mRNA in *Sst*-Cre^TG/TG^ mice was also confirmed by qPCR (Figure 1C). While *Sst*-Cre^+/TG^ mice had slightly reduced *Sst* expression compared to wild type control islets, as has been previously reported (Viollet *et al*. 2017), the difference was not statistically significant. We took advantage of how these *Sst*-Cre^TG/TG^ mice are in effect *Sst*-null mice to test the hypothesis that the absence of SST would decrease the glycemic set point.

**Figure 1.**
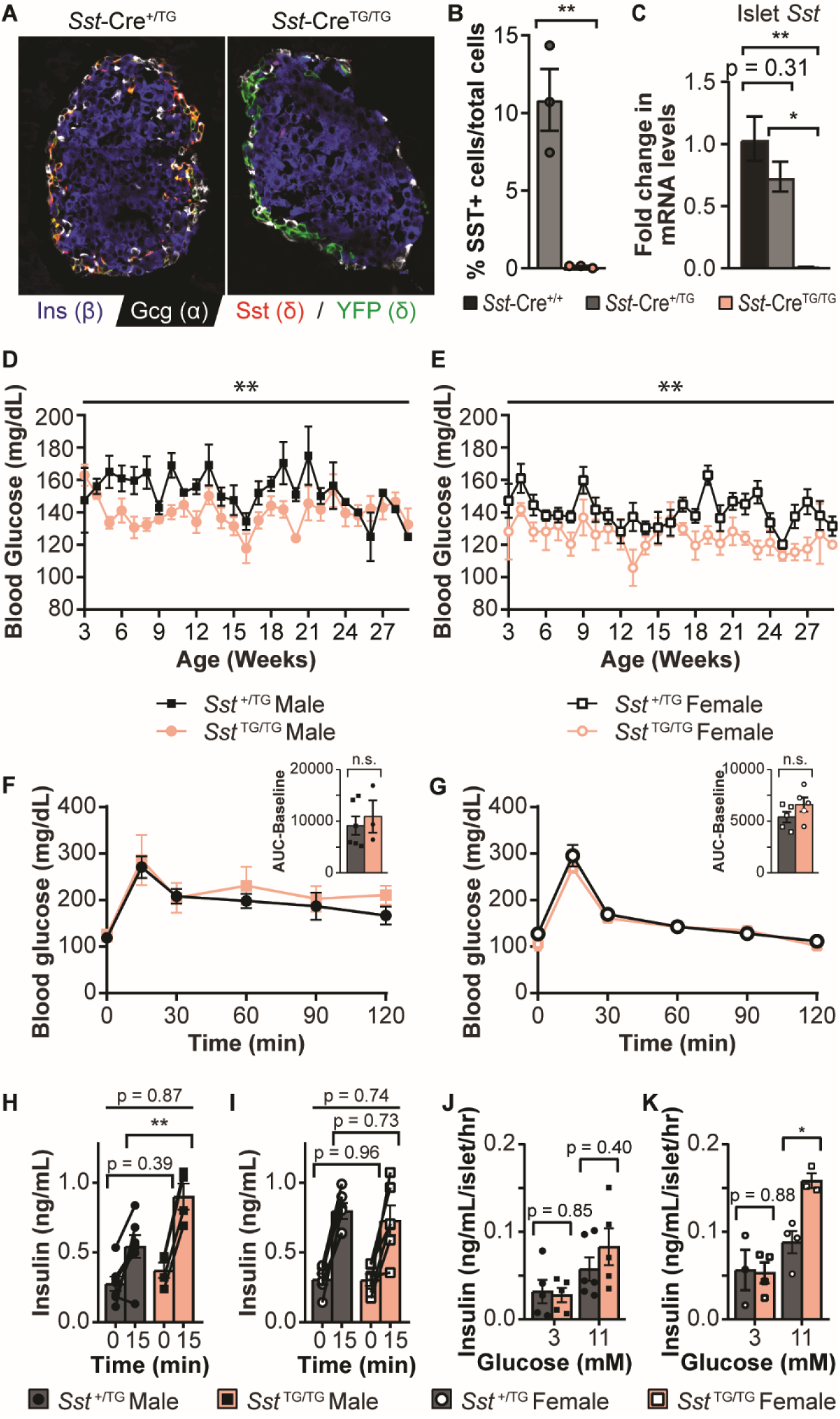
Homozygous *Sst*-Cre mice exhibit loss of *Sst* and a decreased glycemic set point. A) Immunofluorescent stain of pancreas section from a *Sst*-Cre^+/TG^ x lsl-YFP (left) and *Sst*-Cre^TG/TG^ x lsl-YFP mouse (right). Insulin is stained in blue, glucagon in white, SST in red, and YFP in green. Scale bar represents 50 μm. B) Quantification of the number of SST+ cells in *Sst*-Cre^+/TG^ and *Sst*-Cre^TG/TG^ mice (n = 3 mice per group, >10 islets and >500 cells counted per animal). C) *Sst* mRNA levels in *Sst*-Cre^+/+^, *Sst*-Cre^+/TG^, and *Sst*-Cre^TG/TG^ islets. D and E) Weekly blood glucose measurements of (D) male *Sst*-Cre^+/TG^ (n = 6) and *Sst*-Cre^TG/TG^ (n = 9) mice, and E) female *Sst*-Cre^+/TG^ (n = 7) and *Sst*-Cre^TG/TG^ (n = 7) mice, grouped by age. F and G) Glucose tolerance and quantification of the AUC-baseline of male (F) and female (G) mice. H and I) Plasma insulin levels before and 15 min after IP glucose administration in male (H) and female (I) mice. Fold change (insulin levels at 15 minutes divided by insulin levels at 0 minutes), was also compared between *Sst*-Cre^+/TG^ and *Sst*-Cre^TG/TG^ mice. J and K) Static insulin secretion assay using islets from male (J) and female (K) mice incubated at 3 mM glucose and 11 mM glucose. Error bars represent SEM, *p < 0.05, **p < 0.01.

To determine whether the absence of SST affects glycemia, we conducted weekly glucose measurements on the mice. Both male and female *Sst*-Cre^TG/TG^ mice exhibited lower non-fasting glucose levels compared to control *Sst*-Cre^+/TG^ mice of the same sex (Figures 1D and 1E). We then investigated whether there were also changes to fasting and challenged glycemia using an intraperitoneal (IP) glucose tolerance test. Neither sex exhibited significant changes in glucose tolerance (Figures 1F and 1G). While plasma insulin levels were significantly higher in male *Sst*-Cre^TG/TG^ mice relative to male *Sst*-Cre^+/TG^ mice, there was no significant difference in the fold change in insulin levels when we compared plasma insulin levels before and after glucose administration (Figure 1H). There was no significant difference in plasma insulin levels or fold change in insulin between female *Sst*-Cre^+/TG^ and *Sst*-Cre^TG/TG^ mice (Figure 1I). However, islets from female *Sst*-Cre^TG/TG^ mice secreted significantly more insulin under 11 mM glucose (Figure 1J and 1K), matching the phenotype reported for *Sst^-/-^* mice (Hauge-Evans *et al*. 2009).

### δ cell ablation lowers the glycemic set point

Although *Sst*-Cre^TG/TG^ mice have lower non-fasting glycemia and can secrete more insulin in response to the same glucose challenge, the absence of a difference in glucose tolerance suggests that there may be some compensation for the constitutive absence of SST from birth. We therefore turned to a model that would allow us to ablate δ cells and therefore SST signaling within the islet at a time of our choosing using diphtheria toxin (DT). We generated *Sst*-Cre^+/TG^ x Rosa-lsl-iDTR mice (*Sst*-Cre x lsl-DTR mice) expressing the diphtheria toxin receptor (DTR) in SST-expressing cells that would be ablated upon administration of DT. Only *Sst*-Cre^+/TG^ mice were used for these experiments, as *Sst*-Cre^TG/TG^ mice already have lower glycemia (Figure 1). We confirmed complete ablation of δ cells in DT-treated *Sst*-Cre x lsl-DTR mice, while β and α cell numbers did not significantly change relative to saline (SAL)-treated *Sst*-Cre x lsl-DTR littermate controls (Figures 2A and 2B). Similarly, *Sst* mRNA levels were reduced by approximately 40-fold after ablation, while *Ins2* and *Gcg* levels were unaffected (Figure 2C).

**Figure 2.**
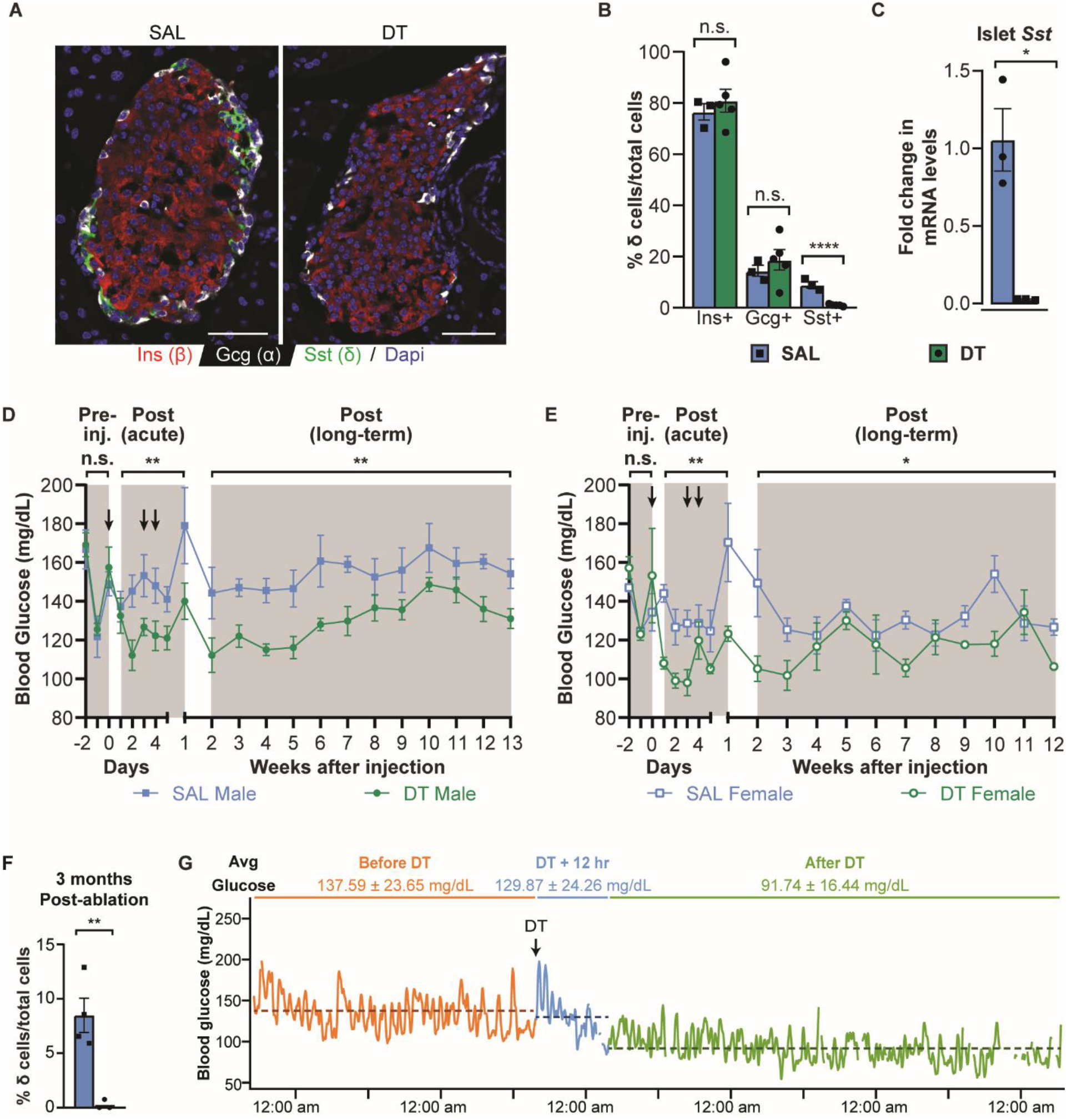
δ cell ablation leads to a lasting decrease in the glycemic set point. A) Immunofluorescent stain of pancreas section from a SAL-treated (left) and DT-treated (right) *Sst*-Cre x lsl-DTR mouse. Insulin is stained in red, glucagon in white, and SST in green. Scale bar represents 50 μm. B) Quantification of the number of insulin, glucagon, and SST+ cells (n = 3 SAL and n = 5 DT mice, >10 islets and >500 cells counted per animal). C) *Sst* mRNA levels in islets from SAL- and DT-treated mice. D and E) Blood glucose measurements of (D) male SAL-treated (n = 4) and DT-treated (n = 6) mice, and E) female SAL-treated (n = 3) and DT-treated (n = 3) mice. Black arrows represent IP administration of SAL or DT. F) Representative CGM data from a mouse. Orange represents glucose levels measured prior to single IP injection of DT. Blue represents the point at which DT was administered and the 12 hours following. Green represents glucose levels measured 12 hours after DT administration until the end of the experiment. Dashed lines represent average glucose level throughout each time period. Error bars represent SEM, *p < 0.05, **p < 0.01, ****p < 0.0001.

SST is also expressed in other tissues throughout the body, primarily the stomach, duodenum, and throughout the hypothalamus. Thus, we collected these tissues to assess the extent of ablation in these areas. Assessing *Sst* mRNA levels by qPCR and staining for SST both revealed that gastric D cells were lost upon acute ablation, but began recovering within two weeks and recovered completely within three months (Figure S1A and S1B). Duodenal D cells were found in mice that experienced ablation and mice that did not, which was reflected by partial ablation immediately following DT administration based on *Sst* transcript level and complete recovery by 2 weeks (Figure S1C and S1D). The at best transient loss of D cells in the gastrointestinal track followed by recovery is in line with the well-established ability of the gastrointestinal mucosae to self-renew within a relatively short period (Barker 2014). In the hypothalamus, there was a slight decrease in *Sst* transcript, while immunofluorescent stains showed that there remained intact SST+ neurons (Figure S1E and S1F). In contrast, pancreatic δ cells do not recover from ablation after 3 months (Figure 2F), consistent with reports that they are long-lived cells without a high turnover rate (Arrojo e Drigo *et al*. 2019).

To determine the effect of δ cell ablation on glycemia, we conducted weekly glucose measurements on the mice, with daily measurements throughout the period of injection. In both sexes, glucose levels between groups were indistinguishable prior to δ cell ablation (Figure 2D and 2E). Following IP injection of DT, both male and female *Sst*-Cre x DTR mice immediately exhibited a significant and lasting decrease in glucose levels compared to *Sst*-Cre x DTR littermates that received SAL. The difference in basal glycemia between the control and δ cell-ablated mice remained even after 3 months. To obtain higher temporal resolution of changes in the glycemic set point, we placed continuous glucose monitors on the mice to obtain glucose profiles with 5-minute resolution as described before (van der Meulen *et al*. 2015). Within 12 hours of a single dose of DT, the blood glucose levels of *Sst*-Cre x DTR mice began to drop and remained steady for the duration of the experiment (Figure 2G). This confirmed that the changes observed through the weekly glucose measurements indeed reflected an acute and lasting change in the glycemic set point of the mice.

We also observed that *Sst*-Cre x lsl-DTR mice treated with DT had a lower body weight relative to controls, although the difference was not significant in female mice (Figure S2A and S2B). This brought up the possibility that lower food intake contributed to the decreased glycemia. It had also been previously demonstrated that ablation of SST-expressing neurons specifically in the tuberal nucleus of the hypothalamus decreases food intake (Luo *et al*. 2018). To determine whether the mice were indeed feeding less, we measured food intake in DT-treated *Sst*-Cre x lsl-DTR mice and *Sst*-Cre littermates without the lsl-DTR transgene. We observed no significant changes in food intake (Figure S2C), suggesting that there is not sufficient ablation within the hypothalamus to affect feeding patterns and that the decrease in glycemia is not due to decreased food intake. Given that the glycemic set point remains lower even after 3 months and only the pancreatic δ cells remain completely absent at that time point, this indicates that the permanent change in the glycemic set point in these mice can be specifically attributed to the ablation of pancreatic δ cells.

### δ cell ablation increases glucose tolerance and glucose-stimulated insulin secretion

To determine the effect of δ cell ablation during glucose stimulation, we conducted IP glucose tolerance tests on the mice before and after administration of SAL or DT. As expected, there was no significant difference between the glucose tolerance of the different groups of mice prior to δ cell ablation (Figures 3A). Glucose tolerance tests performed 36 hours after the last administration of DT revealed that DT-treated male *Sst*-Cre x DTR mice had significantly improved glucose tolerance (Figure 3B). As with males, female *Sst*-cre x DTR mice did not exhibit a significant difference in glucose tolerance prior to injection of SAL or DT (Figure 3C). After injection, DT-treated female mice demonstrated significantly lower fasting blood glucose levels but otherwise no statistically significant difference in overall glucose tolerance, in contrast to our observations in male mice (Figure 3D).

**Figure 3.**
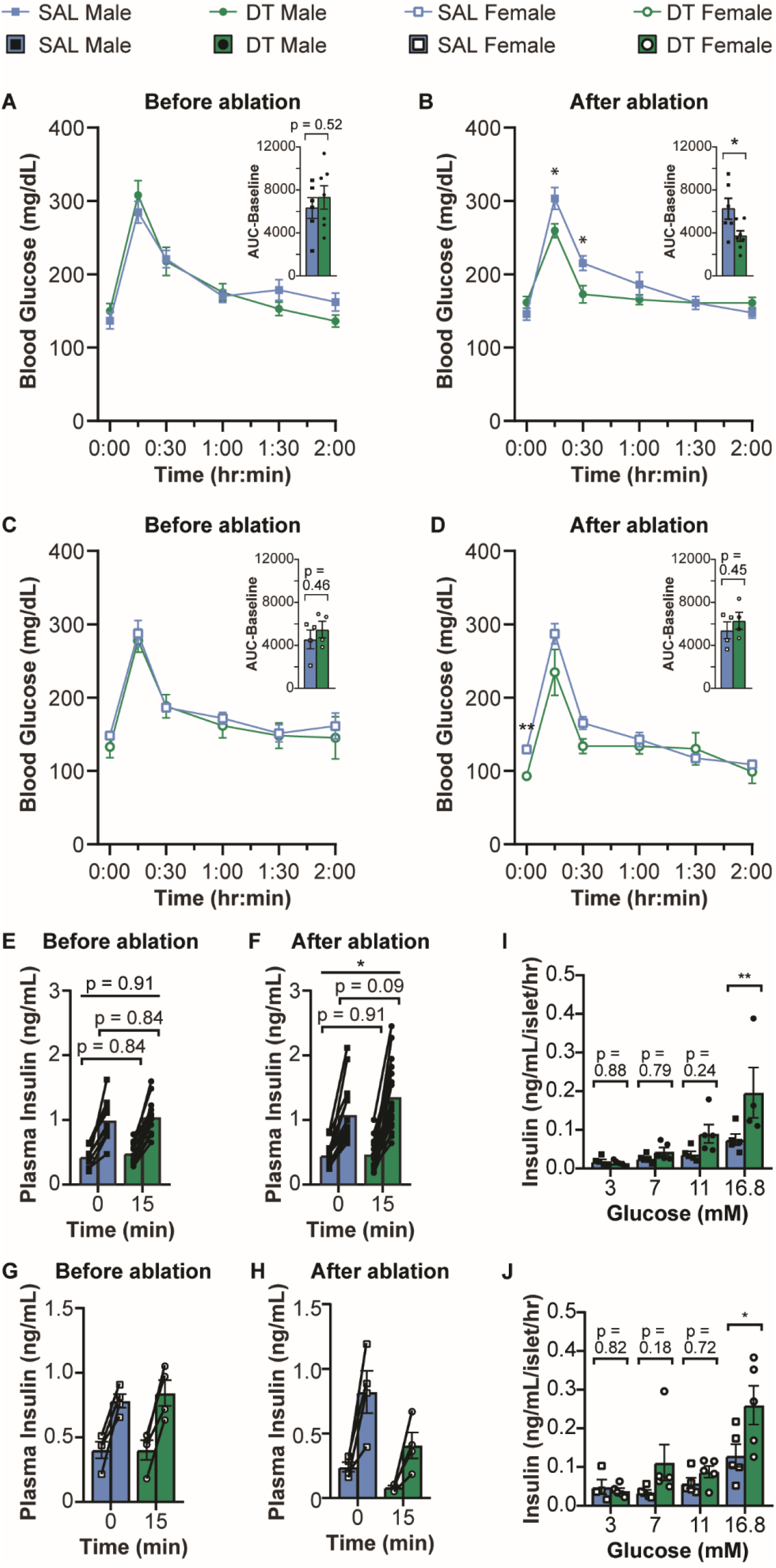
δ cell ablation increases glucose tolerance and insulin secretion. A-D) Glucose tolerance test of male mice A) before δ cell ablation and B) after δ cell ablation (n = 6 SAL, n = 7 DT), and female mice C) before δ cell ablation and D) after δ cell ablation (n = 4 SAL, n = 4 DT). Bar graphs in the upper right-hand corner of each line graph represent AUC-baseline. E-H) Plasma insulin measurements E) male mice before ablation, F) male mice after ablation, G) female mice before ablation, and H) female mice after ablation. Blood for insulin measurement was collected at 0 minutes (before glucose administration) and 15 minutes after glucose administration. The 0 minute time point, 15 minute time point, and fold change in plasma insulin levels are compared. I and J) Static insulin secretion assay performed on islets isolated from ablated I) male and J) female mice. Error bars represent SEM. *p < 0.05, **p < 0.01.

Given that the ablation of δ cells would lead to a decrease in SST tone and remove its local inhibition of β cells, we hypothesized that the decrease in the glycemic set point and increase in glucose tolerance would both be the result of an increase in glucose-stimulated insulin secretion (GSIS). To test this hypothesis, we measured plasma insulin in fasted SAL- or DT-treated *Sst*-Cre x DTR mice before and 15 minutes after IP injection of glucose. Fasting and glucose-stimulated plasma insulin levels were comparable between groups in both sexes prior to ablation (Figure 3E). After δ cell ablation, there was a slight but non-statistically significant increase in glucose-stimulated plasma insulin levels in male (Figure 3F). However, there was a significant increase in the fold-change of plasma insulin after ablation, as determined by dividing plasma insulin levels 15 minutes after glucose injection by baseline plasma insulin levels in each mouse. No differences in plasma insulin levels were observed in females (Figure 3G and 3H).

To confirm our *in vivo* findings *in vitro*, we compared static GSIS in the presence and absence of δ cells in isolated intact islets. This revealed a consistent increase in GSIS at glucose levels mildly or moderately above the β cell glucose threshold, which reached significance at 16.8 mM glucose in islets from both sexes (Figure 3I and 3J). Thus, GSIS increases in the absence of pancreatic δ cells, and the effect is islet autonomous since the increase is observed in isolated islets *in vitro*.

### Paracrine somatostatin secretion from δ cells mediates the changes in the glycemic set point

We next decided to employ the *Sst*-Cre^+/TG^ x *lsl-Gα_i_-DREADD* model (*Sst*-Cre x Gi-DREADD). This allowed us to acutely inhibit δ cell activity and therefore SST secretion with clozapine-N-oxide (CNO) while keeping δ cells intact and for reversal of δ cell inhibition upon removal of CNO. IP administration of CNO (1 mg/kg) led to a decrease in basal blood glucose levels in both male and female *Sst*-Cre x Gi-DREADD mice, as opposed to CNO administered to mice without the Cre driver (Figure 4A and 4B), consistent with the decrease in glucose levels seen in δ cell-ablated mice. 24 hours after CNO administration, there was no significant difference in non-fasting glucose levels between the CNO-treated *Sst*-Cre x Gi-DREADD group (CNO) and SAL-treated *Sst*-Cre x Gi-DREADD or CNO-treated Gi-DREADD only group (CTRL) in either sex. Glucose tolerance tests performed in independent experiments 1 hour after administration of CNO showed that inhibition of δ cell activity also improved glucose tolerance (Figure 4C and 4D). The long-term effect of continuous CNO was assessed by implanting osmotic pumps containing 1 mg/mL CNO dissolved in 0.9% saline or saline alone into *Sst*-Cre x Gi-DREADD females. Blood glucose levels were comparable prior to implantation, while a decrease in blood glucose levels was observed in *Sst*-Cre x Gi-DREADD females that received osmotic pumps with CNO (Figure S3). Blood glucose levels appeared to increase following removal of CNO-filled osmotic pumps, which may possibly be due to increased SST secretion following removal of inhibition. Thus, inhibition of δ cell activity has similar effects to ablation of δ cells.

**Figure 4.**
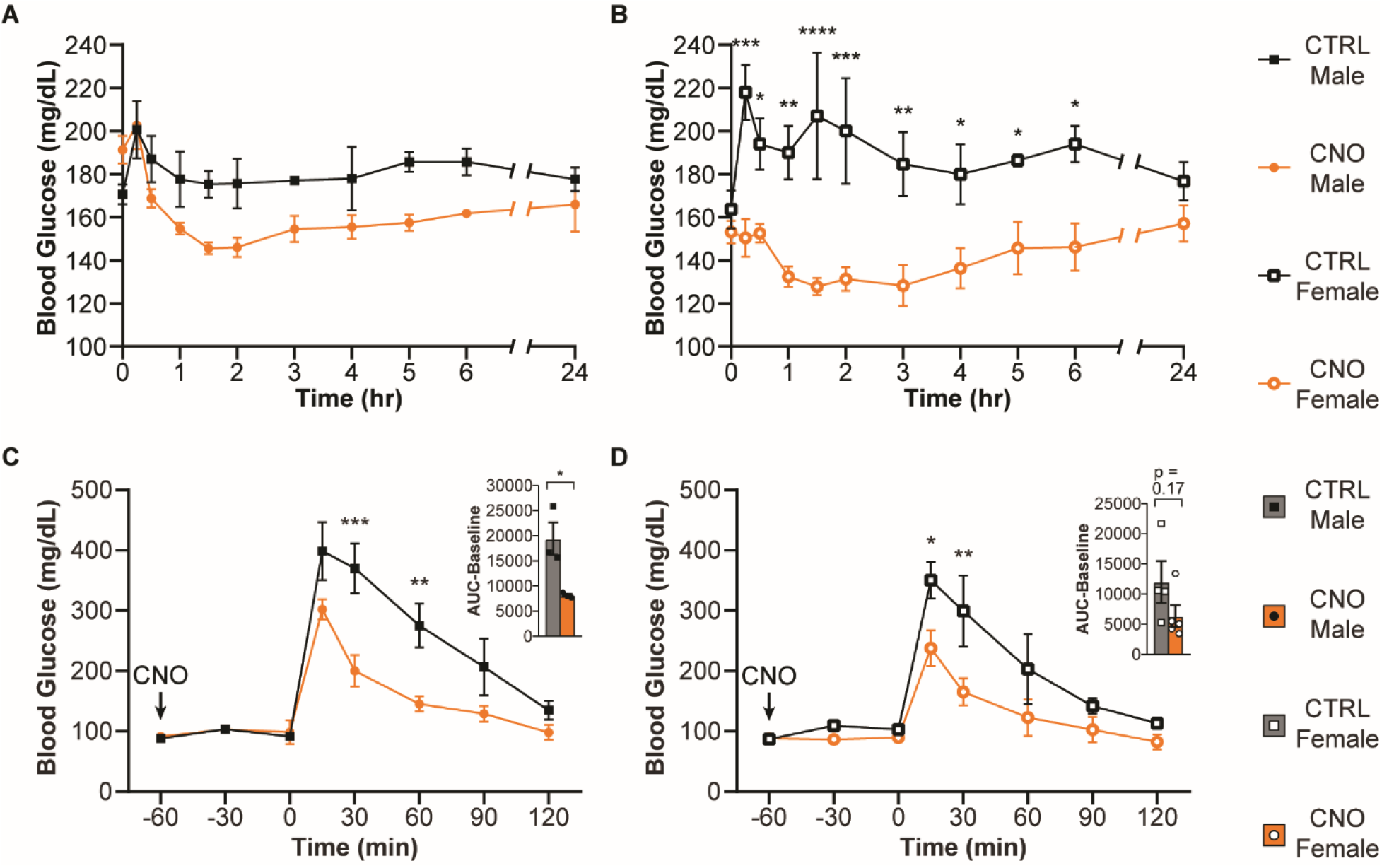
Inhibition of δ cell activity decreases glycemia. A and B) Hourly glucose measurements after administration of 1 mg/kg CNO at t = 0 min in A) male (n = 3 CTRL, n = 4 CNO) and B) female (n = 4 CTRL and n = 5 CNO) mice. Mice were used at 5 months of age. C and D) Glucose tolerance test after CNO administration 1 hour before IP injection of glucose in C) male (n = 3 CTRL, n = 3 CNO) and D) female (n = 4 CTRL, n = 4 CNO) mice. Mice were used at 6 months of age. Bar graphs in the upper right-hand corner of each line graph represent AUC-baseline. Error bars represent SEM. *p < 0.05, **p < 0.01, ***p < 0.001, ****p < 0.0001.

### α cell ablation does not affect basal glycemia

Since SST inhibits glucagon secretion, it is possible that the drop in glycemic set point and increase insulin secretion we observe upon δ cell ablation are indirectly mediated by increased glucagon release. The role of the α cell in amplifying insulin secretion induced by nutrient stimulation in the prandial state is increasingly appreciated. Removing glucagon signaling to the β cell by knocking out glucagon or its receptor, or inhibiting α cell activity using Gi-DREADD all lead to decreased GSIS (Svendsen *et al*. 2018; Capozzi *et al*. 2019a, b; Zhu *et al*. 2019). This demonstrates that although glucagon is generally thought of as a hormone that acts to raise glucose levels, it also potentiates GSIS to bring glucose levels down in the prandial state. Furthermore, a recent comprehensive and elegantly executed paper establishing that the glycemic set point is set by paracrine interactions within the islet concluded that the underlying mechanism was paracrine crosstalk between β and α cells (Rodriguez-Diaz *et al*. 2018). However, mouse models of α cell ablation generally demonstrated no change in glycemia nor insulin secretion (Thorel *et al*. 2011; Pedersen *et al*. 2013; Shiota *et al*. 2013), with the exception of moderately decreased GSIS from a pancreas perfusion (Svendsen *et al*. 2018). Thus, we considered it important to juxtapose the outcomes of α cell ablation to δ cell ablation conducted by the same investigators to directly compare the effect of δ and α cell ablation on glycemia and insulin secretion.

We set up *Gcg*-CreER x lsl-DTR mice to enable DT-mediated ablation of α cells, with the *Gcg*1-CreER line chosen for its superior and more specific labeling of α cells (Ackermann *et al*. 2016). Successful ablation of α cells was confirmed by immunofluorescence analysis and qPCR (Figure 5A-C), as with the ablation of δ cells. α cell ablation did not have a significant effect on the glycemic set point in mice of both sexes (Fig 5D and 5E). There was also no significant difference in glucose tolerance and plasma insulin levels between groups of mice both before and after administration of DT (Figure 5F-M). These findings are in close agreement with previous studies that investigated changes in glycemia after α cell ablation (Thorel *et al*. 2011; Pedersen *et al*. 2013; Shiota *et al*. 2013). Therefore, our findings confirm that ablating α cells in mice does not alter non-fasting blood glucose levels, in sharp contrast to our experiments where we ablated δ cells using the same methodology.

**Figure 5.**
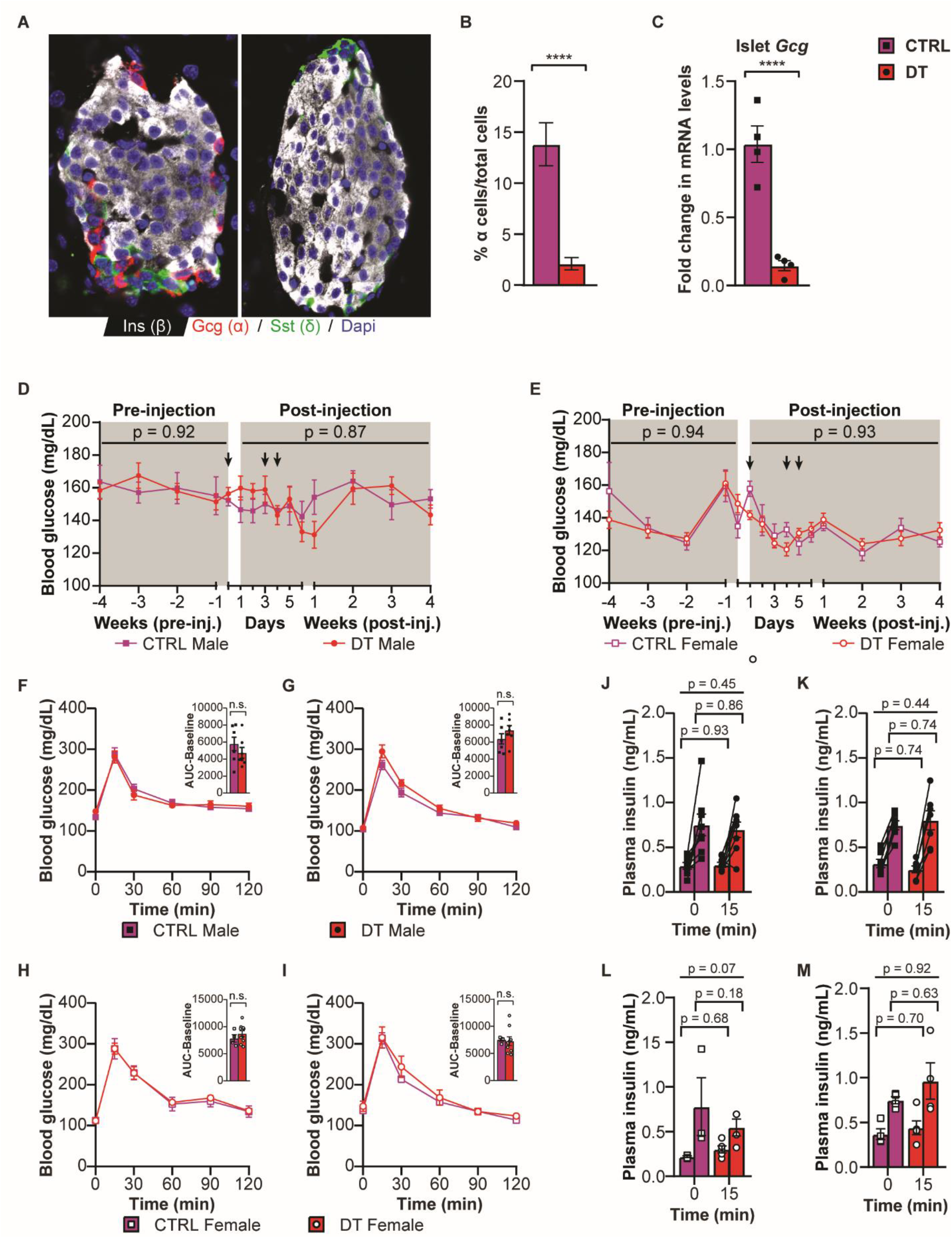
α cell ablation does not affect basal glycemia. A) Immunofluorescent stain of pancreas section from a non-ablated (left) and α cell-ablated mouse (right). Insulin is stained in white, glucagon in red, and SST in green. Scale bar represents 50 μm. B) Quantification of α cell number. C) *Gcg* mRNA levels in control and ablated mice. D and E) Glucose measurements of D) male (n = 5 CTRL, 5 DT) and E) female mice (n = 4 CTRL, 10 DT). Arrows represent IP administration of DT. F-I) Glucose tolerance tests in male mice F) before α cell ablation and G) after α cell ablation, and female mice H) before α cell ablation, and I) after α cell ablation. Bar graphs in the upper right corner of each line graph represent AUC-baseline. J-M) Plasma insulin levels in J) male mice before ablation, K) male mice after ablation, L) female mice before ablation, and M) female mice after ablation.

To determine whether absence of δ cells leads to increased α cell activity, we stimulated α cells with 100 nM epinephrine under low glucose. Epinephrine-induced glucagon secretion significantly increased in islets without δ cells, suggesting that loss of paracrine SST from δ cell ablation does indeed disinhibit α cells (Figure S4A). This is in line with other studies that have established the role of δ cells in restraining α cell glucagon release upon prandial stimulation (Lai *et al*. 2018; Xu *et al*. 2020; Singh *et al*. 2021). To determine whether glucagon signaling contributes to the increase in insulin secretion observed at different glucose concentrations, we also measured glucagon concentration from the same samples. A general pattern of higher glucagon secretion in the absence of δ cells was observed, with significantly higher glucagon secretion from islets from male mice that were incubated in 16.8 mM glucose (Figure S4B and S4C). This suggests that the absence of local SST signaling leads to reduced inhibition of α cell activity under high glucose, and this may contribute to the increase GSIS observed at higher glucose levels well above the glucose threshold for insulin secretion.

### δ cell ablation decreases the glucose threshold for β cell response

To more precisely assess the effect of the absence of δ cells on β cell activity in response to glucose at the single cell level, we turned to calcium imaging. As calcium is necessary for insulin secretion, changes in intracellular calcium levels are an excellent proxy for insulin secretion. Importantly, calcium imaging using genetically-encoded sensors such as GCaMP6s allows for single-cell resolution and subsequent fixation and *post hoc* immunofluorescent staining to validate the identity of the recorded cells. To this end, we generated quadruple transgenic mice expressing *MIP*-Cre/ERT x *Sst*-Cre x lsl-DTR x lslGCaMP6. In this line, mice constitutively express both DTR and GCaMP6 in SST-expressing cells. After δ cell ablation with DT, tamoxifen administration to the mice allows for the translocation of Cre/ERT to the nucleus, activating GCaMP6 expression in insulin-expressing β cells. The simultaneous induction of DTR in β cells does not lead to β cell ablation as DT is no longer administered. This strategy allows us to ablate δ cells, then observe β cell calcium response. Due to the complexity of the cross, we used a mix of SAL-treated *MIP*-Cre/ERT x *Sst*-cre^+/TG^ x DTR x GCaMP6 mice and DT-treated mice expressing *MIP*-Cre/ERT x GCaMP6 with or without DTR or *Sst*-cre as controls. There is no significant difference between *Sst*-Cre only, lsl-DTR only, or *Sst*-Cre x lsl-DTR mice prior to administration of DT (Figure S5). We hypothesized based on our observations that the loss of δ cells would shift the β cell glucose threshold to the left. To test this hypothesis, we performed glucose step experiments starting below the β cell glucose threshold at 4 mM glucose and increasing in 1 mM increments every 20 minutes, observing islets with and without δ cells simultaneously in the same microfluidic chamber. After each trace, islets were fixed to confirm the absence of δ cells in ablated islets and to ensure that each GCaMP6 expressing cell was insulin-positive via *post hoc* whole mount immunofluorescence.

For analysis of β cell calcium response, we defined the activity threshold as the half-max of the signal for each individual β cell. We then determined the glucose level at which each cell first reaches that threshold of activity. In control islets with intact δ cells, individual β cells generally began responding shortly after exposure to 6 or 7 mM glucose, with a synchronized response between the majority of β cells by 8 or 9 mM glucose (Figure 6A, Video S1) and presence of δ cells confirmed by a *post hoc* stain (Figure 6B). In islets with ablated δ cells, individual β cells generally began to respond shortly after exposure to 5 or 6 mM glucose, with a synchronized response by 7 or 8 mM glucose (Figure 6C, Video S1) and absence of δ cells confirmed by a *post hoc* stain (Figure 6D). Quantified across over 2000 β cells from at least 10 islets per mouse in 3 pairs of mice, the β cell glucose threshold in islets without δ cells was on average 1 mM glucose (approximately 18 mg/dL) lower and statistically significantly different from islets with intact δ cell feedback (Figure 6E and 6F).

**Figure 6.**
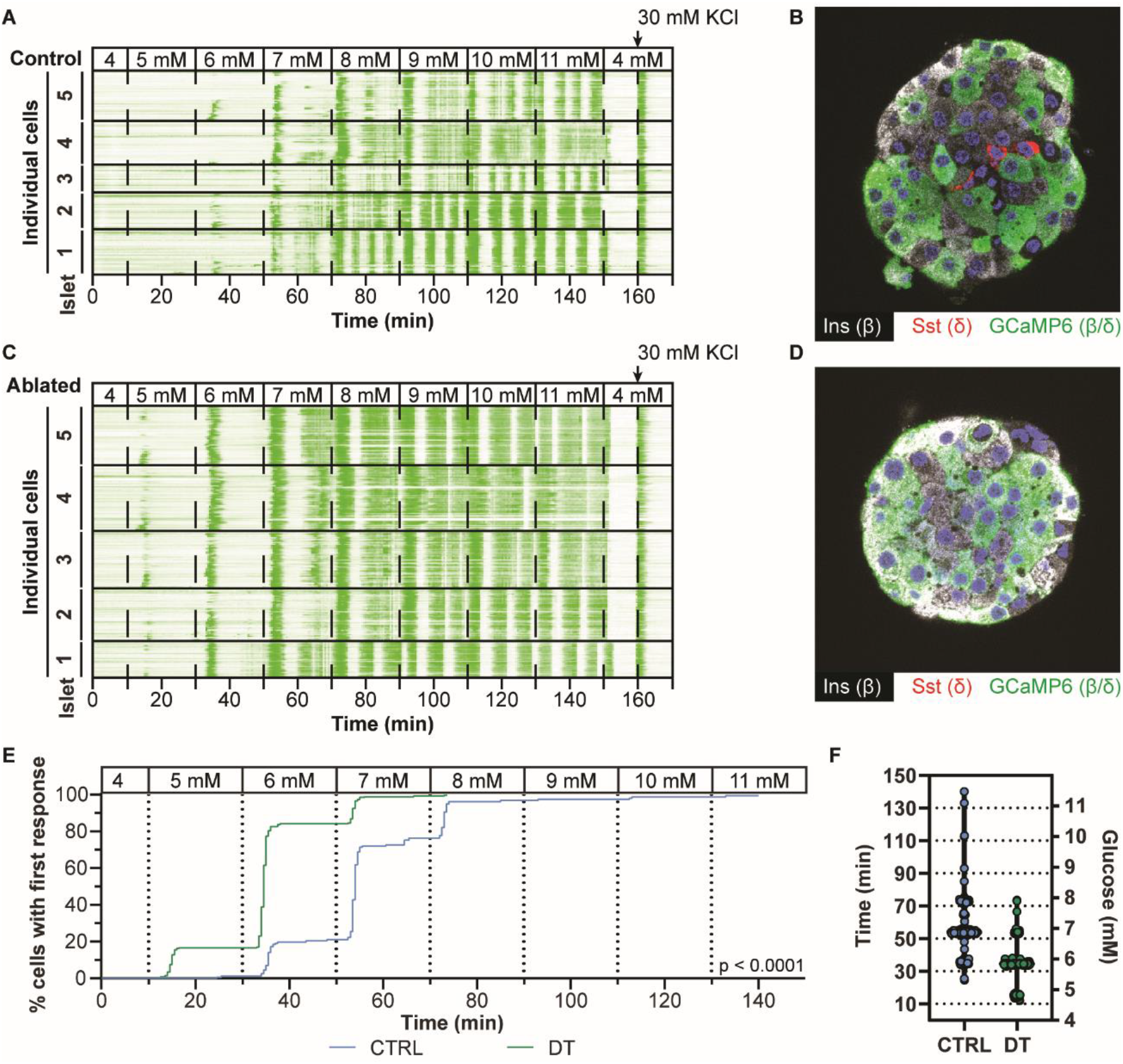
β cells exhibit calcium response at a lower glucose threshold in the absence of δ cells. A) Representative trace of β cells from a mouse with intact pancreatic δ cells. Each line represents the calcium activity of a single β cell, with green corresponding to an increase in intracellular calcium. Each box represents an islet. Dashed lines represent the point at which the glucose levels were changed. 30 mM KCl was added at the end of each trace to confirm viability of the cells. B) *Post hoc* whole mount stain of an islet with intact δ cells. Scale bar represents 50 μm. C) Representative trace of β cells from a mouse with ablated δ cells. D) *Post hoc* whole mount stain of an islet confirming absence of δ cells. E) Curve representing the % of cells that have their first response at each glucose level, with the first response defined as the point at which a cell first reaches half-max of its signal intensity. F) Violin plot in which each dot represents a cell and the point at which it first responded.

To determine whether the shift in calcium response also represents a shift in insulin secretion, we simultaneously imaged β cell calcium activity while collecting the outflow for measurement of insulin secretion. This demonstrated that β cells secrete insulin at a higher amplitude in islets without δ cells (Figure 7). From these experiments, we concluded that the ablation of δ cells decreases the glucose threshold at which β cells become active and observe that the 1 mM, or 18 mg/dL decrease in the glucose threshold is similar to the approximately 20-30 mg/dL decrease in the glycemic set point observed in mice *in vivo*.

**Figure 7.**
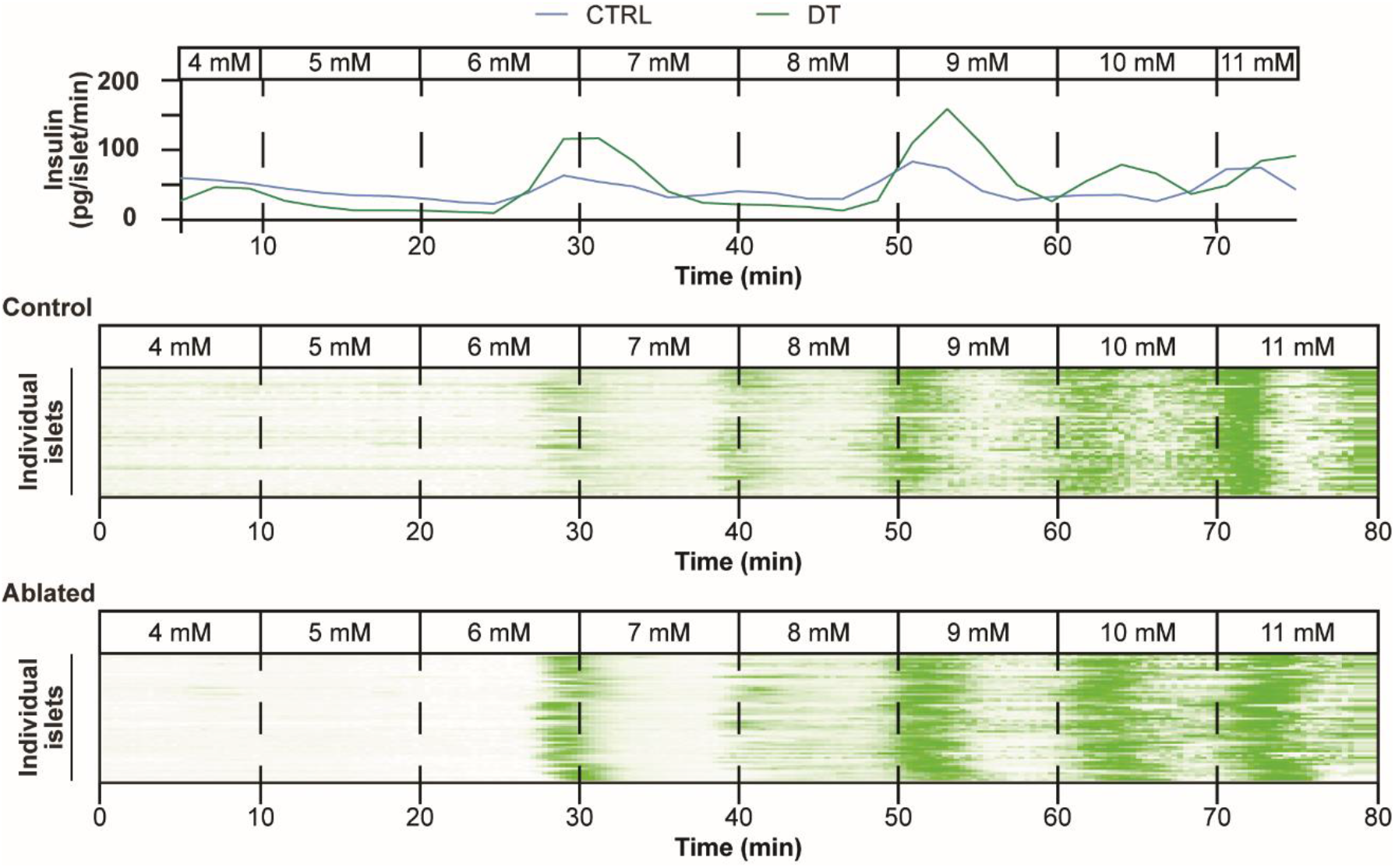
Simultaneous collection of calcium dynamics in islets and insulin secretion. Secretion dynamics are on the top. Below are calcium traces of whole islets from control or δ cell-ablated mice.

## DISCUSSION

SST has long been known to inhibit both β and α cells (Brazeau *et al*. 1973; Koerker *et al*. 1974; Dubois 1975). Secretion of SST from δ cells has been proposed to prevent excess insulin secretion (van der Meulen *et al*. 2015; Huising *et al*. 2018) and has been demonstrated to inhibit glucagon secretion in the prandial phase (Lai *et al*. 2018; Xu *et al*. 2020). However, the physiological contribution of local δ cell-mediated feedback on the glycemic set point has not been addressed to date. Here we demonstrate that δ cells determine the glycemic set point in mice by modulating the glucose threshold of pancreatic β cells. Both male and female mice exhibit an immediate and sustained decrease in basal blood glucose levels upon δ cell ablation. Male mice also exhibit an increase in glucose tolerance that occurs due to an increase in the fold-change of plasma insulin secreted in response to glucose. Static secretion assays of isolated islets confirmed an increase in GSIS from islets in both sexes. This suggests that there may be other physiological factors in female mice that prevent the increase in plasma insulin levels seen in male mice; for example, females are protected from systemic insulin resistance compared to males (Hevener *et al*. 2018; De Paoli *et al*. 2021), which may explain why they did not exhibit changes in glucose tolerance. Calcium imaging with single-cell resolution revealed that in the absence of δ cells, β cells respond to lower levels of glucose, demonstrating increased glucose sensitivity. The consistency with which β cells respond at a 1 mM lower glucose level in islets without δ cells is in line with the approximately 1 mM decrease in blood glucose levels. Furthermore, we used *Sst*-Cre x Gi-DREADD mice to show that inhibition of δ cell activity is sufficient to decrease glycemia and increase glucose tolerance. Altogether, these data demonstrate that removing paracrine SST secretion from δ cells within the pancreatic islet leads to a decreased glycemic set point.

One *Sst*-null mouse model was previously observed to have increased non-fasting blood glucose levels at 3 weeks of age (Richardson *et al*. 2015). Another *Sst*-null mouse model has also been noted to have lower blood glucose levels but no differences in non-fasting insulin levels (Luque & Kineman 2007). These mice also display no differences in glucose tolerance relative to wild-type mice but a slight elevation in fasting insulin (Gahete *et al*. 2011; Luque *et al*. 2016), matching our observations with *Sst*-Cre^TG/TG^ mice. Thus, it is indeed likely that there are compensatory effects when SST is constitutively absent.

Due to SST having many effects throughout the body, including but not limited to the brain, the stomach, and the intestines, we took care to determine their potential contributions. The duodenum is unlikely to contribute since there is little ablation and any ablation that does occur recovers within 2 weeks. SST in the gut has also been implicated to promote satiety (Lieverse *et al*. 1995). The stomach is also unlikely to contribute since it recovers within 3 months, at which point the mice still exhibit a decreased glycemic set point. Furthermore, D cells in the stomach inhibit gastric secretion by inhibiting gastrin release (Bloom *et al*. 1974; Holst *et al*. 1992), and suppressing gastric motility (Johansson *et al*. 1981) and gastric emptying (Holst *et al*. 2016). Collectively, the effect of D cells is to decrease the nutrient absorption rate. D cell ablation would therefore be predicted to instead increase circulating nutrients, at least post-prandially. Therefore, the observed reduction of blood glucose upon δ cell ablation or inhibition is inconsistent with reduced D cell function, but consistent with reduced δ cell-mediated inhibition of β cells.

Since the hypothalamus plays an important role in energy metabolism, it is possible that ablation of SST-expressing cells there would contribute to the decrease in glycemia. SST neurons have already been implicated in food and water intake (Karasawa *et al*. 2014; Stengel & Taché 2019). While SST-expressing neurons in the tuberal nucleus promote food intake (Luo *et al*. 2018), we ruled out potentially confounding effects of hypothalamic SST neuron ablation by observing no changes in food intake between mice with and without δ cells. Our observations are consistent with a recent paper reporting that specifically ablating SST-neurons in the hypothalamus by stereotaxic injection of DTA, the catalytic unit of diphtheria toxin, had no effect on blood glucose or non-fasting insulin levels (Huang *et al*. 2021). Collectively, this suggests that the consistent reductions in the glycemic set point we observe upon DT administration is not due to ablation of SST-expressing neurons.

A recent paper reported that ablation of pancreatic δ cells leads to neonatal death, and the authors attributed it to hypoglycemia (Li *et al*. 2018). In contrast, the mice used in our study remained healthy months after ablation and demonstrate a moderate and stable reduction in their blood glucose. This difference is likely due to their use of the lsl-DTA mouse (Ivanova *et al*. 2005), which expresses the catalytic subunit of diphtheria toxin constitutively from the onset of *Sst*-dependent expression of Cre. This would cause immediate and continued ablation of all SST-expressing cells, including those in the CNS, and likely lead to developmental CNS defects such as motor neuron defects that may interfere with the neonatal pups’ ability to feed. This could lead to failure to thrive and feed normally and may explain the observed hypoglycemia in this specific model. The conclusion that the absence of pancreatic δ cells leads to neonatal death is inconsistent with multiple mouse models, including our δ cell ablation model, multiple *Sst*-null mice (Luque & Kineman 2007; Hauge-Evans *et al*. 2009; Viollet *et al*. 2017), and also *Hhex*-KO mice that do not develop pancreatic δ cells due to absence of the essential transcription factor HHEX (Zhang *et al*. 2014). These mice are all viable, making it unlikely that pancreatic δ cell ablation is primarily responsible for the early perinatal lethality of *Sst*-Cre x lsl-DTA mice. While our data agree that pancreatic δ cell ablation leads to increased insulin secretion and decreased glucose levels, the viability of our mice allowed us to demonstrate that the hypoglycemia is not transient but a stable change in the glycemic set point of these mice. This also illustrates the utility of the DTR-mediated ablation model over the DTA mouse line; our DTR model allows us to choose the timing of ablation and dose the DT to allow for selective ablation of peripheral cells without also ablating SST-expressing neurons.

Our data agrees with previous studies that ablation of α cells does not have an effect on glycemia in mice as demonstrated by two previous studies (Thorel *et al*. 2011; Pedersen *et al*. 2013). Thus, while α cells undoubtedly play an important role in both the counterregulatory response through systemic action and the stimulation of insulin secretion through paracrine action in the prandial phase, their contribution appears to be dispensable around the non-fasting glycemic set point in mice. However, in human islets the paracrine interactions and islet interactions are substantially different (Steiner *et al*. 2010; Caicedo 2013; Noguchi & Huising 2019; Walker *et al*. 2021). α cells may therefore contribute more directly to the glycemic set point in human islets (Rodriguez-Diaz *et al*. 2018).

Since these experiments were all performed in the context of healthy mice, it would be interesting to observe the effect that the absence of δ cells can have in models of diabetes, and whether they would ameliorate the progression due to the subsequent increase in glucose sensitivity of β cells, or exacerbate it due to β cells reaching a state of exhaustion more quickly. Whether δ cells play a similar role in humans, who have a lower glycemic set point, is an important question to be resolved in subsequent studies as well.

In summary, our findings illustrate that pancreatic δ cells determine the glycemic set point through their interaction with β cells. Our findings establish the physiological role of δ cells as local dampeners of insulin secretion during the resting state. Upon removal or inhibition of δ cells, we observe an immediate lowering of the glycemic set point. These observations are also consistent with the notable increase in the glycemic set point known to occur around 2-3 weeks postnatally in mice (Rozzo *et al*. 2009; Blum *et al*. 2012) that we demonstrated is caused by onset of β cell UCN3 expression and subsequent increase of SST tone (van der Meulen *et al*. 2015). Our findings quantify the contribution of pancreatic δ cells and simultaneously indicate that their physiological role is to restrain β cells and moderate insulin secretion, but not absolutely prevent nutrient stimulation of β cell insulin secretion. As such, δ cells are a prime example of the important and complementary role that paracrine feedback inhibition plays in determining important physiological parameters such as our glycemic set point. δ cell feedback accomplishes this in concert with the β cell autonomous glucose threshold, acting as redundant mechanisms that collectively safeguard against inappropriate and acutely dangerous hyperinsulinemic hypoglycemia.

## ACKNOWLEDGEMENTS

This work was supported by the National Institute of Diabetes and Digestive and Kidney Disease (NID DK-110276). J.L.H was supported by a National Institute of General Medical Sciences-funded Pharmacology Training Program (T32 GM-099608). S.L. was supported by the NSF Graduate Research Fellowship (1650042), the UC Davis Training Program in Molecular and Cellular Biology (T32 GM-007377), and the UC Davis NSF Bridge to Doctorate Program (1612490). M.S.P. is supported by the UC Davis Training Program in Molecular and Cellular Biology (T32 GM-007377).

## AUTHOR CONTRIBUTIONS

Conceptualization, J.L.H. and M.O.H.; Methodology, J.L.H. S.L., and M.O.H.; Software, M.S.P.; Validation, J.L.H., S.L., and M.S.P.; Formal Analysis, J.L.H., S.L., and M.S.P.; Investigation, J.L.H., S.L., M.S.P., N.K., J.V.G., N.C., P.A., and S.K.; Writing - Original Draft, J.L.H and M.O.H; Visualization, J.L.H. and M.O.H.; Supervision, M.O.H.

## METHODS

### Animals

Mice were maintained in group housing (4 mice per cage) on a 12 hr light:12 hr darkness cycle with water and standard rodent chow provided *ad libitum*. Heterozygous *Sst*-IRES-Cre mice (*Sst^tm2.1(cre)Zjh^*/J, Jax #013044) (Taniguchi *et al*. 2011) were crossed together to generate homozygous *Sst*-IRES-Cre mice (*Sst*-Cre^TG/TG^). *Sst*-Cre^+/TG^ x lsl-YFP (B6.129X1-*Gt(ROSA)26Sor^tm1(EYFP)Cos^*/J, Jax # 006148) (Srinivas *et al*. 2001) mice were crossed to *Sst*-Cre^+/TG^ to generate heterozygous and homozygous *Sst*-Cre x lsl-YFP mice. *Sst*-Cre x lsl-DTR mice were initially generated by crossing *Sst*-Cre^+/TG^ mice to homozygous *R26-iDTR* mice (C57BL/6-*Gt(ROSA)26Sor^tm1(HBEGF)Awai^*/J, Jax # 007900) (Buch *et al*. 2005), then maintained by crossing bi-transgenic offspring to C57BL/6N mice or by crossing mice with complementary transgenes. For tdTomato lineage-labeling, some of the mice were also crossed to *Ai14(RCL-tdT)-D* mice (B6.Cg-*Gt(ROSA)26Sor^tm14(CAG-tdTomato)Hze/J^*, Jax # 007914) (Madisen *et al*. 2010). To ablate alpha cells, mice expressing *Gcg*-CreERT2 (B6;129S4-*Gcg^em1(cre/ERT2)Khk^*/Mmjax, Jax #030346) (Ackermann *et al*. 2016) were also crossed to lsl-DTR mice. For beta cell calcium imaging, *Sst*-Cre x lsl-DTR mice were crossed to mice expressing *MIP-CreERT* (B6.Cg-Tg(Ins1-cre/ERT)1Lphi/J, Jax # 024709) (Wicksteed *et al*. 2010) and *Ai96(RCL-GCaMP6s)* (B6;129S6-*Gt(ROSA)26 Sor^tm96(CAG-GCaMP6s)Hze^*/J, Jax # 24106) (Madisen *et al*. 2015), then maintained by crossing mice expressing complementary transgenes, with one parent expressing lsl-DTR and the other expressing lsl-GCaMP6. *Sst*-Cre x lsl-Gi-DREADD mice were generated by crossing *Sst*-Cre^+/TG^ mice to homozygous *R26-LSL-Gi-DREADD* mice (B6.129-Gt(ROSA)26 *Sor^tm1(CAG-CHRM4*,-mCitrine)Ute^*/J, Jax # 026219) (Zhu *et al*. 2016), then maintained by crossing bi-transgenic offspring to C57BL/6N mice or to complementary littermates. *Sst*-Cre^TG/TG^ mice were not used for breeding. Mice were used between 2 and 4 months of age unless otherwise indicated. All mouse experiments were approved by the UC Davis Institutional Animals Care and Use Committee and were performed in compliance with the Animal Welfare Act and the Institute for Laboratory Animal Research Guide to the Care and Use of Laboratory Animals.

### DT, tamoxifen, and CNO treatments

For delta cell ablation, 126 ng diphtheria toxin (List Biological Laboratories, Product # 150) in 200 μL 0.9% saline was injected into mice via IP injection on days 0, 3, and 4. The same timeline for DT administration was followed for alpha cell ablation, except 300 ng was given. Control mice were given an intraperitoneal injection on the same days with an equivalent volume of 0.9% saline. Tamoxifen (Sigma-Aldrich, Catalog # T5648) was dissolved in sunflower oil (Trader Joe’s, Monrovia, CA, USA) at 20 mg/mL, then administered to mice via oral gavage with a volume of 250 μL for 5 consecutive days. CNO (Tocris Catalog # 4936) was dissolved in 0.9% saline and administered to mice at 1 mg/kg via IP injection.

### Glucose tolerance test and plasma insulin collection

Mice were fasted overnight for 16 hours. The next morning, they were weighed and put into individual cages. Tails were clipped with a surgical scissor and baseline glucose measured before administration of 2 mg/kg glucose via IP injection (Dextrose, Sigma-Aldritch, Catalog # D9559). *Sst*-Cre x Gi-DREADD mice were administered 1 mg/kg CNO via IP injection 1 hour before glucose. All blood glucose measurements over the 2-3 hour time period were done using a OneTouch Ultra glucometer. To collect plasma insulin, tail blood from mice was collected using the Microvette CB300 EDTA (Sarstedt, Product # 16.444.100) and kept on ice. After all the 15 min time points were collected, the samples were spun down at 4°C at 5000 rpm for 10 minutes and the plasma collected into a non-stick tube (Ambion, Catalog # AM12300). Samples were stored at −20°C until assayed.

### Islet isolation

Islets were isolated by injecting 2 mL collagenaseP (0.8 mg/mL in HBSS, Roche Diagnostics, Catalog # 11249002001) into the bile duct with the ampulla of Vater clamped. The pancreas was removed into a conical tube to which an additional 2 mL of collagenaseP was added, then incubated at 37°C for 11 min. This was followed by gentle manual shaking to dissociate the pancreata, then three washes with cold HBSS + 5% NCS (Newborn Calf Serum). After the digested suspension was passed through a nylon mesh (pore size 425 μm, Small Parts), the islets were isolated by density gradient centrifugation using Histopaque (Sigma-Aldrich, Catalog # 10771) for 20 min at 1400 xg without brake. Islets were then collected from the interface, washed with cold HBSS + 5% NCS, and hand-picked several times under a dissecting microscope, followed by culture in RPMI + 5.5 mM Glucose + 10% FBS + pen/strep.

### Static insulin secretion assays

Islets were picked twice into Krebs Ringer Buffer (20 mM Hepes pH 7.4, 1.2 mM KH_2_PO_4_, 25 mM NaHCO_3_, 130 mM NaCl, 5 mM KCl, 1.2 mM MgCl_2_, 1.2 mM CaCl_2_) containing 0.1% BSA and 3 mM glucose, then incubated at 37°C for 1 hour. For each group, islets were pooled and split into different treatments with at least 5 replicates each. Static insulin secretion was carried out using 10 islets per well, with different treatments added after the islets had been placed into the wells.

### Calcium imaging and dynamic insulin secretion assays

Microfluidics chambers were bonded to 35 mm dishes with a glass bottom (Mattek, # P35G-1.5-14-C). Islets were set down into these chambers and allowed to adhere to the glass by overnight culture. Continuous perfusion of Krebs Ringer Buffer at a rate of 200 μL per minute was maintained using the Elveflow microfluidics system, with different treatments adjusted using the Mux distributor. The calcium response of islets over time was imaged using a Nikon Eclipse Ti2 using a 60x lens with oil. Simultaneous collection of dynamic insulin perfusate was done by collecting the outflow into non-stick tubes.

### Hormone measurements

Plasma insulin was measured using the Ultra-Sensitive Mouse Insulin ELISA Kit wide range assay (Crystal Chem, Catalog # 90080). Secreted insulin was measured using the Insulin LUMIT kit (Promega, # CS3037A01) in 384-well plates (Corning, #3572) at 10 μL (static secretion) or 25 μL (dynamic secretion) sample volumes.

### Immunofluorescence and cell counting

Pancreata were isolated with the spleen and fixed in 4% PFA for 5 hours, then protected with 30% sucrose overnight prior to embedding with OCT (Fisher Healthcare, Catalog # 4585). Cryosections of 14 μM thickness were collected using a Leica cryostat. The same procedure was conducted with the stomach, which was isolated, halved lengthwise, and washed twice in PBS prior to fixation in PFA, and the duodenum, which was collected as an approximately 1 cm piece adjacent to the stomach and opened up prior to fixation. The brain was fixed in 4% borate PFA for 24 hours, protected with 30% sucrose overnight, and collected as 50 μM thick sections using a vibratome. For immunofluorescence, slides were first washed for 5 minutes three times in KBS, then incubated with antibodies diluted in donkey block (KPBS supplemented with 2% donkey serum and 0.4% Triton X-100) overnight at 4°C. Tissue samples were incubated with secondary antibodies diluted in donkey block the following day and mounted using ProLong Gold Antifade Mountant (Invitrogen, # P36930). Slides were imaged on a Nikon Eclipse Ti using a 60x lens with oil. For whole mount staining of islets after calcium imaging, islets were fixed in 4% PFA in the chamber for 15 minutes at room temperature, then washed in PBS twice before a 15-minute incubation at 4°C. Islets were incubated in donkey block overnight at 4°C, followed by overnight incubation with primary antibodies diluted in donkey block, overnight wash in PBS + 0.15% Tween 20, and overnight incubation with secondary antibodies diluted in donkey block. Finally, islets were incubated in 4% PFA either overnight at 4°C or for 1 hour at room temperature, followed by several washes in PBS + 0.15% Tween 20 every 30 minutes. After the washes, the islets were put in Rapiclear (SunJin Lab, Catalog # RC152001) and imaged on a Nikon Eclipse Ti2. Cells were counted manually using ImageJ.

### RNA Extraction and qPCR

Tissues were collected into TRIzol Reagent (Invitrogen, #15596026). Prior to the start of the RNA extraction, hypothalamus, stomach, and duodenal tissue collected into TRIzol were sonicated. Islets were directly broken down in TRIzol. RNA was isolated by chloroform extraction and precipitated by isopropanol. Once pellets were resuspended, cDNA was made using a High-Capacity cDNA Reverse Transcription Kit (Applied Biosystems, Catalog # 4368813). qPCR was performed using the PowerUp SYBR Green Master Mix (Applied Biosystems, Catalog # A25741) or iTaq Universal SYBR Green Supermix (Bio-Rad, Catalog # 1725121) on a Bio-Rad CFX 384.

### Continuous Glucose Monitoring

Mice were anesthetized, shaved, and sterilized. A Dexcom G6 sensor was introduced subcutaneously into mice and bonded using veterinary glue. Receivers were left adjacent to cages and monitored. Continuous Glucose Monitoring profiles were collected for approximately a week.

### Osmotic Pump Implantation

For osmotic pump delivery of CNO, we used the ALZET Osmotic Pump Model 2001, which has a nominal flow rate of 1.0 μL/hr for 1 week. The pumps were filled with approximately 200 μL CNO (1 mg/mL) in 0.9% saline, or 0.9% saline alone. After the mice were anesthetized, shaved, and sterilized, the pump was inserted into a subcutaneous pocket made with an incision to the dorsal base of the neck.

### Statistical Analysis

Data were analyzed by Student’s t-test, corrected for multiple comparisons using the Holm-Sidak method where appropriate, and represented as mean ± SEM, with n representing number of animals in each group. Differences were considered significant when p < 0.05. Statistics were computed using Prism (GraphPad Software, La Jolla, CA).

## REFERENCES

Ackermann AM, Zhang J, Heller A, Briker A & Kaestner KH 2016 High-fidelity Glucagon-CreER mouse line generated by CRISPR-Cas9 assisted gene targeting. Molecular Metabolism 6 236–244. (doi:10.1016/j.molmet.2017.01.003)

Alonge KM, D’Alessio DA & Schwartz MW 2021 Brain control of blood glucose levels: implications for the pathogenesis of type 2 diabetes. Diabetologia 64 5–14. (doi:10.1007/s00125-020-05293-3)

Arrojo e Drigo R, Jacob S, García-Prieto CF, Zheng X, Fukuda M, Nhu HTT, Stelmashenko O, Peçanha FLM, Rodriguez-Diaz R, Bushong E et al. 2019 Structural basis for delta cell paracrine regulation in pancreatic islets. Nature Communications 10 3700. (doi:10.1038/s41467-019-11517-x)

Barker N 2014 Adult intestinal stem cells: Critical drivers of epithelial homeostasis and regeneration. Nature Reviews Molecular Cell Biology 15 19–33. (doi:10.1038/nrm3721)

Bloom SR, Mortimer CH, Thorner MO, Besser GM, Hall R, Gomez-Pan A, Roy VM, Russell RCG, Coy DH, Kastin AT et al. 1974 Inhibition of Gastrin and Gastric-Acid Secretion By Growth-Hormone Release-Inhibiting Hormone. The Lancet 304 1106–1109. (doi:10.1016/S0140-6736(74)90869-1)

Blum B, Hrvatin S, Schuetz C, Bonal C, Rezania A & Melton DA 2012 Functional beta-cell maturation is marked by an increased glucose threshold and by expression of urocortin 3. Nature Biotechnology 30 261–264. (doi:10.1038/nbt.2141)

Blum B, Roose AN, Barrandon O, Maehr R, Arvanites AC, Davidow LS, Davis JC, Peterson QP, Rubin LL & Melton DA 2014 Reversal of β cell de-differentiation by a small molecule inhibitor of the TGFβ pathway. ELife 3 e02809. (doi:10.7554/eLife.02809)

Brazeau P, Vale W, Burgus R, Ling N, Butcher M, Riviver J & Guillemin R 1973 Hypothalamic polypeptide that inhibits the secretion of immunoreactive pituitary growth hormone. Science 179 77–79.

Buch T, Heppner FL, Tertilt C, Heinen TJAJ, Kremer M, Wunderlich FT, Jung S & Waisman A 2005 A Cre-inducible diphtheria toxin receptor mediates cell lineage ablation after toxin administration. Nature Methods 2 419–426. (doi:10.1038/nmeth762)

Caicedo A 2013 Paracrine and autocrine interactions in the human islet: More than meets the eye. Seminars in Cell and Developmental Biology 24 11–21. (doi:10.1016/j.semcdb.2012.09.007)

Capozzi ME, Wait JB, Koech J, Gordon AN, Coch RW, Svendsen B, Finan B, D’Alessio DA & Campbell JE 2019a Glucagon lowers glycemia when β cells are active. JCI Insight 4. (doi:10.1172/jci.insight.129954)

Capozzi ME, Svendsen B, Encisco SE, Lewandowski SL, Martin MD, Lin H, Jaffe JL, Coch RW, Haldeman JM, MacDonald PE et al. 2019b β Cell tone is defined by proglucagon peptides through cAMP signaling. JCIInsight 4. (doi:10.1172/jci.insight.126742)

Carroll PB, Zeng Y, Alejandro R, Starzl TE & Ricordi C 1992 Glucose homeostasis is regulated by donor islets in xenografts. Transplantation Proceedings 24 2980–2981.

Chambers AP, Sorrell JE, Haller A, Roelofs K, Hutch CR, Kim KS, Gutierrez-Aguilar R, Li B, Drucker DJ, D’Alessio DA et al. 2017 The Role of Pancreatic Preproglucagon in Glucose Homeostasis in Mice. Cell Metabolism 25 927–934.e3. (doi:10.1016/j.cmet.2017.02.008)

Dubois MP 1975 Immunoreactive somatostatin is present in discrete cells of the endocrine pancreas. Proceedings of the National Academy of Sciences of the United States of America 72 1340–1343. (doi:10.1073/pnas.72.4.1340)

Ewing KL & Tauber OE 1964 Blood chemistry changes in mice fed high levels of polyoxyethylene sorbitan derivatives. Toxicology and Applied Pharmacology 6 442–446. (doi:10.1016/S0041-008X(64)80010-7)

Gahete MD, Pozo-salas AI, Martínez-fuentes AJ, Lecea L De, Gracia-navarro F, Kineman RD, Castan JP & Luque RM 2011 Cortistatin Is Not a Somatostatin Analogue but Stimulates Prolactin Release and Inhibits GH and ACTH in a Gender-Dependent Fashion : Potential Role of Ghrelin. 152 4800–4812. (doi:10.1210/en.2011-1542)

Gerich JE 1993 Control of Glycaemia. (doi:10.1016/S0950-351X(05)80207-1)

German JP, Thaler JP, Wisse BE, Oh-I S, Sarruf DA, Matsen ME, Fischer JD, Taborsky GJ, Schwartz MW & Morton GJ 2011 Leptin activates a novel CNS mechanism for insulin-independent normalization of severe diabetic hyperglycemia. Endocrinology 152 394–404. (doi:10.1210/en.2010-0890)

Hauge-Evans AC, King AJ, Carmignac D, Richardson CC, Robinson ICAF, Low MJ, Christie MR, Persaud SJ & Jones PM 2009 Somatostatin secreted by islet δ-cells fulfills multiple roles as a paracrine regulator of islet function. Diabetes 58 403–411. (doi:10.2337/db08-0792)

Hevener AL, Zhou Z, Moore TM, Drew BG & Ribas V 2018 The impact of ERα action on muscle metabolism and insulin sensitivity – Strong enough for a man, made for a woman. Molecular Metabolism 15 20–34. (doi:10.1016/j.molmet.2018.06.013)

Holst JJ, Orskov C & Seier-Poulsen S 1992 Somatostatin is an essential paracrine link in acid inhibition of gastrin secretion. Digestion 51 95–102. (doi:10.1159/000200882)

Holst JJ, Gribble F, Horowitz M & Rayner CK 2016 Roles of the gut in glucose homeostasis. Diabetes Care 39 884–892. (doi:10.2337/dc16-0351)

Huang C, Rosencrans RF, Bugescu R, Vieira CP, Hu P, Adu-Agyeiwaah Y, Gamble KL, Longhini ALF, Fuller PM, Leinninger GM et al. 2021 Depleting hypothalamic somatostatinergic neurons recapitulates diabetic phenotypes in mouse brain, bone marrow, adipose and retina. Diabetologia 64 2575–2588. (doi:10.1007/s00125-021-05549-6)

Huising MO 2020 Paracrine regulation of insulin secretion. Diabetologia 63 2057–2063. (doi:10.1007/s00125-020-05213-5)

Huising MO, van der Meulen T, Huang JL, Pourhosseinzadeh MS & Noguchi GM 2018 The Difference δ-Cells Make in Glucose Control. Physiology 33 403–411. (doi:10.1152/physiol.00029.2018)

Ivanova A, Signore M, Caro N, Greene NDE, Copp AJ & Martinez-Barbera JP 2005 In vivo genetic ablation by Cre-mediated expression of diphtheria toxin fragment A. Genesis 43 129–135. (doi:10.1002/gene.20162)

Johansson C, Wisen O, Efendic S & Uvnäs-Wallensten K 1981 Effects of Somatostatin on Gastrointestinal Propagation and Absorption of Oral Glucose in Man. Digestion 22 126–137. (doi:https://doi.org/10.1159/000198619)

Karasawa H, Yakabi S, Wang L, Stengel A, Rivier J & Taché Y 2014 Brain somatostatin receptor 2 mediates the dipsogenic effect of central somatostatin and cortistatin in rats: Role in drinking behavior. American Journal of Physiology - Regulatory Integrative and Comparative Physiology 307 R793–R801. (doi:10.1152/ajpregu.00248.2014)

Kavalakatt S, Khadir A, Madhu D, Hammad M, Devarajan S, Abubaker J, Al-Mulla F, Tuomilehto J & Tiss A 2019 Urocortin 3 Levels Are Impaired in Overweight Humans With and Without Type 2 Diabetes and Modulated by Exercise. Frontiers in Endocrinology 10 1–11. (doi:10.3389/fendo.2019.00762)

Koerker DJ, Ruch W, Chideckel E, Palmer J, Goodner CJ, Ensinck J & Gale CC 1974 Somatostatin: Hypothalamic inhibitor of the endocrine pancreas. Science 184 482–484. (doi:10.1126/science.184.4135.482)

Lai BK, Chae H, Gómez-Ruiz A, Cheng P, Gallo P, Antoine N, Beauloye C, Jonas JC, Seghers V, Seino S et al. 2018 Somatostatin is only partly required for the glucagonostatic effect of glucose but is necessary for the glucagonostatic effect of KATP channel blockers. In Diabetes, pp 2239–2253. (doi:10.2337/db17-0880)

Li N, Yang Z, Li Q, Yu Z, Chen X, Li JC, Li B, Ning SL, Cui M, Sun JP et al. 2018 Ablation of somatostatin cells leads to impaired pancreatic islet function and neonatal death in rodents article. Cell Death and Disease 9. (doi:10.1038/s41419-018-0741-4)

Lieverse RJ, Jansen JBMJ, Masclee AAM & Lamers CBHW 1995 Effects of somatostatin on human satiety. Neuroendocrinology 61 112–116. (doi:10.1159/000126831)

Liu L, Dattaroy D, Simpson KF, Barella LF, Cui Y, Xiong Y, Jin J, König GM, Kostenis E, Roman JC et al. 2021 Gq signaling in α cells is critical for maintaining euglycemia. JCI Insight 6 1–17. (doi:10.1172/jci.insight.152852)

Luo SX, Huang J, Li Q, Mohammad H, Lee CY, Krishna K, Kok AMY, Tan YL, Lim JY, Li H et al. 2018 Regulation of feeding by somatostatin neurons in the tuberal nucleus. Science 361 76–81. (doi:10.1126/science.aar4983)

Luque RM & Kineman RD 2007 Gender-Dependent Role of Endogenous Somatostatin in Regulating Growth Hormone-Axis Function in Mice. 148 5998–6006. (doi:10.1210/en.2007-0946)

Luque RM, Cordoba-chacon J & Pozo-salas AI 2016 Obesity-and gender-dependent role of endogenous somatostatin and cortistatin in the regulation of endocrine and metabolic homeostasis in mice. Nature Publishing Group 1–12. (doi:10.1038/srep37992)

Madisen L, Zwingman TA, Sunkin SM, Oh SW, Zariwala HA, Gu H, Ng LL, Palmiter RD, Hawrylycz MJ, Jones AR et al. 2010 A robust and high-throughput Cre reporting and characterization system for the whole mouse brain. Nature Neuroscience 13 133–140. (doi:10.1038/nn.2467)

Madisen L, Garner AR, Shimaoka D, Chuong AS, Klapoetke NC, Li L, van der Bourg A, Niino Y, Egolf L, Monetti C et al. 2015 Transgenic mice for intersectional targeting of neural sensors and effectors with high specificity and performance. Neuron 85 942–958. (doi:10.1016/j.neuron.2015.02.022)

Matschinsky FM & Davis EA 1998 The Distinction between ‘Glucose Setpoint’, ‘Glucose Threshold’ and ‘Glucose Sensor’ is Critical for Understanding the Role of the Pancreatic β-Cell in Glucose Homeostasis. Molecular and Cell Biology of Type 2 Diabetes and Its Complications 14 14–29. (doi:10.1159/000060872)

van der Meulen T, Xie R, Kelly OG, Vale WW, Sander M & Huising MO 2012 Urocortin 3 Marks Mature Human Primary and Embryonic Stem Cell-Derived Pancreatic Alpha and Beta Cells. PLoS ONE 7 1–12. (doi:10.1371/journal.pone.0052181)

van der Meulen T, Donaldson CJ, Cáceres E, Hunter AE, Cowing-Zitron C, Pound LD, Adams MW, Zembrzycki A, Grove KL & Huising MO 2015 Urocortin3 mediates somatostatin-dependent negative feedback control of insulin secretion. Nature Medicine 21 769–776. (doi:10.1038/nm.3872)

Morton GJ, Matsen ME, Bracy DP, Meek TH, Nguyen HT, Stefanovski D, Bergman RN, Wasserman DH & Schwartz MW 2013 FGF19 action in the brain induces insulin-independent glucose lowering. Journal of Clinical Investigation 123 4799–4808. (doi:10.1172/JCI70710)

Noguchi GM & Huising MO 2019 Integrating the inputs that shape pancreatic islet hormone release. Nature Metabolism 1 1189–1201. (doi:10.1038/s42255-019-0148-2)

Pagliara AS, Stillings SN, Hover B, Martin DM & Matschinsky FM 1974 Glucose modulation of amino acid induced glucagon and insulin release in the isolated perfused rat pancreas. Journal of Clinical Investigation 54 819–832. (doi:10.1172/JCI107822)

De Paoli M, Zakharia A & Werstuck GH 2021 The Role of Estrogen in Insulin Resistance: A Review of Clinical and Preclinical Data. American Journal of Pathology 191 1490–1498. (doi:10.1016/j.ajpath.2021.05.011)

Pedersen J, Ugleholdt RK, Jørgensen SM, Windeløv JA, Grunddal K V, Schwartz TW, Füchtbauer EM, Poulsen SS, Holst PJ & Holst JJ 2013 Glucose metabolism is altered after loss of L cells and α-cells but not influenced by loss of K cells. American Journal of Physiology - Endocrinology and Metabolism 304 60–73. (doi:10.1152/ajpendo.00547.2011)

Richardson CC, To K, Foot VL, Hauge-Evans AC, Carmignac D & Christie MR 2015 Increased perinatal remodelling of the pancreas in somatostatin-deficient mice: Potential role of transforming growth factor-beta signalling in regulating beta cell growth in early life. Hormone and Metabolic Research 47 56–63. (doi:10.1055/s-0034-1390427)

Rodriguez-Diaz R, Molano RD, Weitz JR, Abdulreda MH, Berman DM, Leibiger B, Leibiger IB, Kenyon NS, Ricordi C, Pileggi A et al. 2018 Paracrine Interactions within the Pancreatic Islet Determine the Glycemic Set Point. Cell Metabolism 27 549–558.e4. (doi:10.1016/j.cmet.2018.01.015)

Rozzo A, Meneghel-Rozzo T, Delakorda SL, Yang SB & Rupnik M 2009 Exocytosis of insulin: In vivo maturation of mouse endocrine pancreas. In Annals of the New York Academy of Sciences, pp 53–62. (doi:10.1111/j.1749-6632.2008.04003.x)

Ryan KK, Kohli R, Gutierrez-Aguilar R, Gaitonde SG, Woods SC & Seeley RJ 2013 Fibroblast growth factor-19 action in the brain reduces food intake and body weight and improves glucose tolerance in male rats. Endocrinology 154 9–15. (doi:10.1210/en.2012-1891)

Shiota C, Prasadan K, Guo P, El-Gohary Y, Wiersch J, Xiao X, Esni F & Gittes GK 2013 α-cells are dispensable in postnatal morphogenesis and maturation of mouse pancreatic islets. American Journal of Physiology - Endocrinology and Metabolism 305 1030–1040. (doi:10.1152/ajpendo.00022.2013)

Singh B, Khattab F, Chae H, Desmet L, Herrera PL & Gilon P 2021 KATP channel blockers control glucagon secretion by distinct mechanisms: A direct stimulation of α-cells involving a [Ca2+]c rise and an indirect inhibition mediated by somatostatin. Molecular Metabolism 53 101268. (doi:10.1016/j.molmet.2021.101268)

Srinivas S, Watanabe T, Lin CS, William CM, Tanabe Y, Jessell TM & Costantini F 2001 Cre reporter strains produced by targeted insertion of EYFP and ECFP into the ROSA26 locus. BMC Developmental Biology 1 1–8. (doi:10.1186/1471-213X-1-4)

Steiner DJ, Kim A, Miller K & Hara M 2010 Pancreatic islet plasticity: Interspecies comparison of islet architecture and composition. Islets 2 135–145. (doi:10.4161/isl.2.3.11815)

Stengel A & Taché Y 2019 Central somatostatin signaling and regulation of food intake. Annals of the New York Academy of Sciences 1455 98–104. (doi:10.1111/nyas.14178)

Svendsen B, Larsen O, Gabe MBN, Christiansen CB, Rosenkilde MM, Drucker DJ & Holst JJ 2018 Insulin Secretion Depends on Intra-islet Glucagon Signaling. Cell Reports 25 1127–1134.e2. (doi:10.1016/j.celrep.2018.10.018)

Taniguchi H, He M, Wu P, Kim S, Paik R, Sugino K, Kvitsani D, Fu Y, Lu J, Lin Y et al. 2011 A Resource of Cre Driver Lines for Genetic Targeting of GABAergic Neurons in Cerebral Cortex. Neuron 71 995–1013. (doi:10.1016/j.neuron.2011.07.026)

Tellez K, Hang Y, Gu X, Chang CA, Stein RW & Kim SK 2020 In vivo studies of glucagon secretion by human islets transplanted in mice. Nature Metabolism 2 547–557. (doi:10.1038/s42255-020-0213-x)

Thorel F, Damond N, Chera S, Wiederkehr A, Thorens B, Meda P, Wollheim CB & Herrera PL 2011 Normal glucagon signaling and β-cell function after near-total α-cell ablation in adult mice. Diabetes 60 2872–2882. (doi:10.2337/db11-0876)

Thorens B 2012 Sensing of glucose in the brain. Handbook of Experimental Pharmacology 209 277–294. (doi:10.1007/978-3-642-24716-3_12)

Vieira E, Salehi A & Gylfe E 2007 Glucose inhibits glucagon secretion by a direct effect on mouse pancreatic alpha cells. Diabetologia 50 370–379. (doi:10.1007/s00125-006-0511-1)

Viollet C, Simon A, Tolle V, Labarthe A, Grouselle D, Loe-Mie Y, Simonneau M, Martel G & Epelbaum J 2017 Somatostatin-IRES-cre mice: Between knockout and wild-type? Frontiers in Endocrinology 8 1–8. (doi:10.3389/fendo.2017.00131)

Walker JT, Saunders DiC, Brissova M & Powers AC 2021 The Human Islet: Mini-Organ with Mega-Impact. Endocrine Reviews 42 605–657. (doi:10.1210/endrev/bnab010)

Wicksteed B, Brissova M, Yan W, Opland DM, Plank JL, Reinert RB, Dickson LM, Tamarina NA, Philipson LH, Shostak A et al. 2010 Conditional gene targeting in mouse pancreatic β-cells: Analysis of ectopic cre transgene expression in the brain. Diabetes 59 3090–3098. (doi:10.2337/db10-0624)

Xu SFS, Andersen DB, Izarzugaza JMG, Kuhre RE & Holst JJ 2020 In the rat pancreas, somatostatin tonically inhibits glucagon secretion and is required for glucose-induced inhibition of glucagon secretion. Acta Physiologica. (doi:10.1111/apha.13464)

Zhang J, Mckenna LB, Bogue CW & Kaestner KH 2014 The diabetes gene Hhex maintains δ - cell differentiation and islet function. Genes and Development 829–834. (doi:10.1101/gad.235499.113)

Zhu H, Aryal DK, Olsen RHJ, Urban DJ, Swearingen A, Forbes S, Roth BL & Hochgeschwender U 2016 Cre-dependent DREADD (Designer Receptors Exclusively Activated by Designer Drugs) mice. Genesis 54 439–446. (doi:10.1002/dvg.22949)

Zhu L, Dattaroy D, Pham J, Wang L, Barella LF, Cui Y, Wilkins KJ, Roth BL, Hochgeschwender U, Matschinsky FM et al. 2019 Intraislet glucagon signaling is critical for maintaining glucose homeostasis. JCI Insight 4. (doi:10.1172/jci.insight.127994)

