## Supplementary Figures for "Paracrine signaling by pancreatic δ cells determines the glycemic set point in mice"

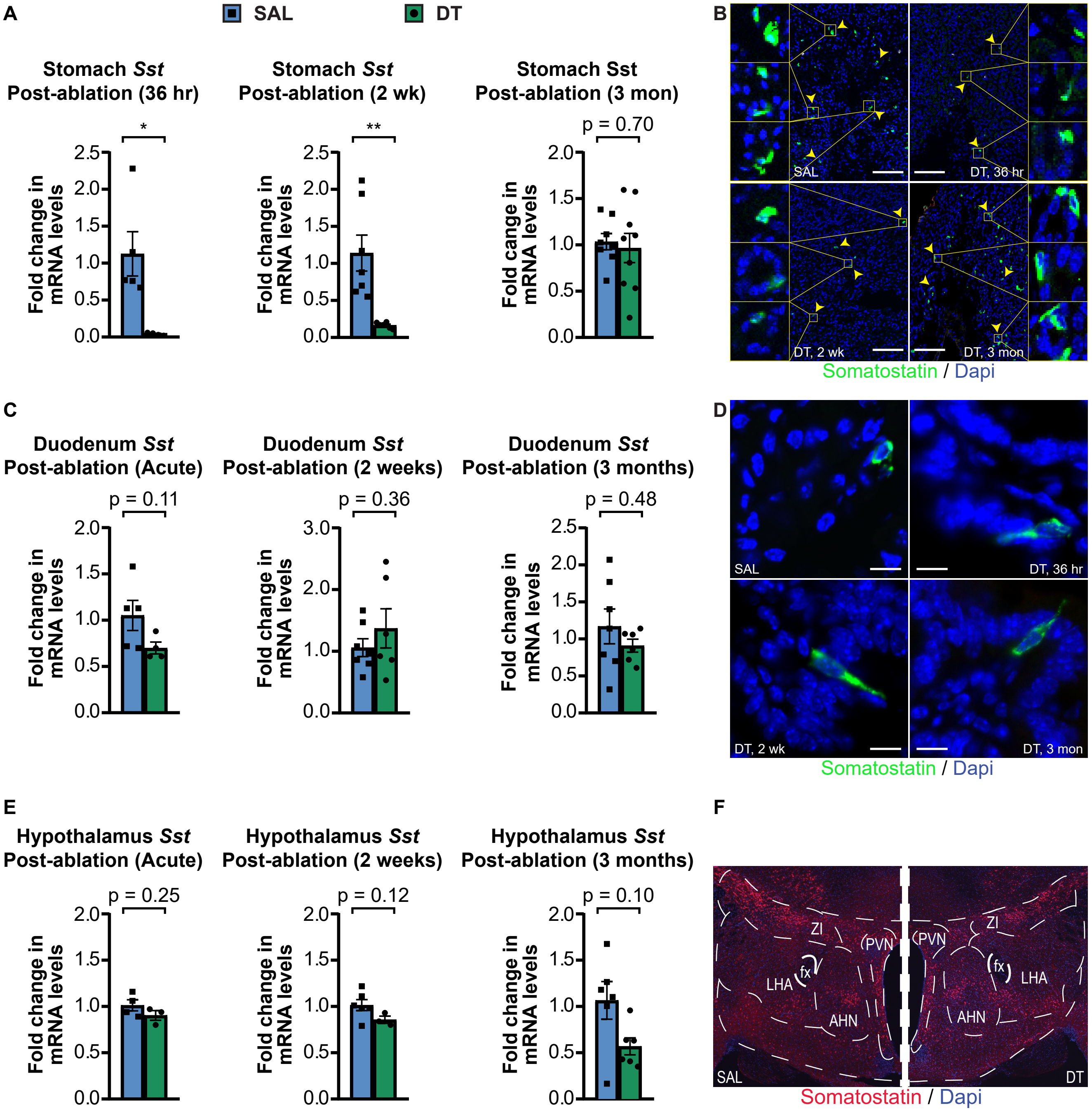


**Figure S1. Transient and partial ablation in other SST-expressing tissues, related to Figure 2.** A) *Sst* mRNA levels in stomach tissue collected from *Sst*-Cre x lsl-DTR mice 36 hours, 2 weeks, or 3 months after administration of SAL or DT. B) SST stain in stomach collected from SAL-treated *Sst*-Cre x lsl-DTR mice and *Sst*-Cre x lsl-DTR mice 36 hours, 2 weeks, or 3 months after DT administration. Yellow arrows indicate gastric D cells and yellow boxes indicate close-ups. Scale bar represents 500 µM. C) *Sst* mRNA levels in duodenum tissue collected from *Sst*-Cre x lsl-DTR mice 36 hours, 2 weeks, or 3 months after administration of SAL or DT. D) SST stain in duodenum collected from SAL-treated *Sst*-Cre x lsl-DTR mice and *Sst*-Cre x lsl-DTR mice 36 hours, 2 weeks, or 3 months after DT administration. Scale bar represents 50 µM. E) *Sst* mRNA levels in hypothalamus collected from *Sst*-Cre x lsl-DTR mice 36 hours, 2 weeks, or 3 months after administration of SAL or DT. F) SST stain in hypothalamus collected from *Sst*-Cre x lsl-DTR mice 6 months after SAL or DT treatment. Error bars represent SEM, *$p < 0.05$, **$p < 0.01$.

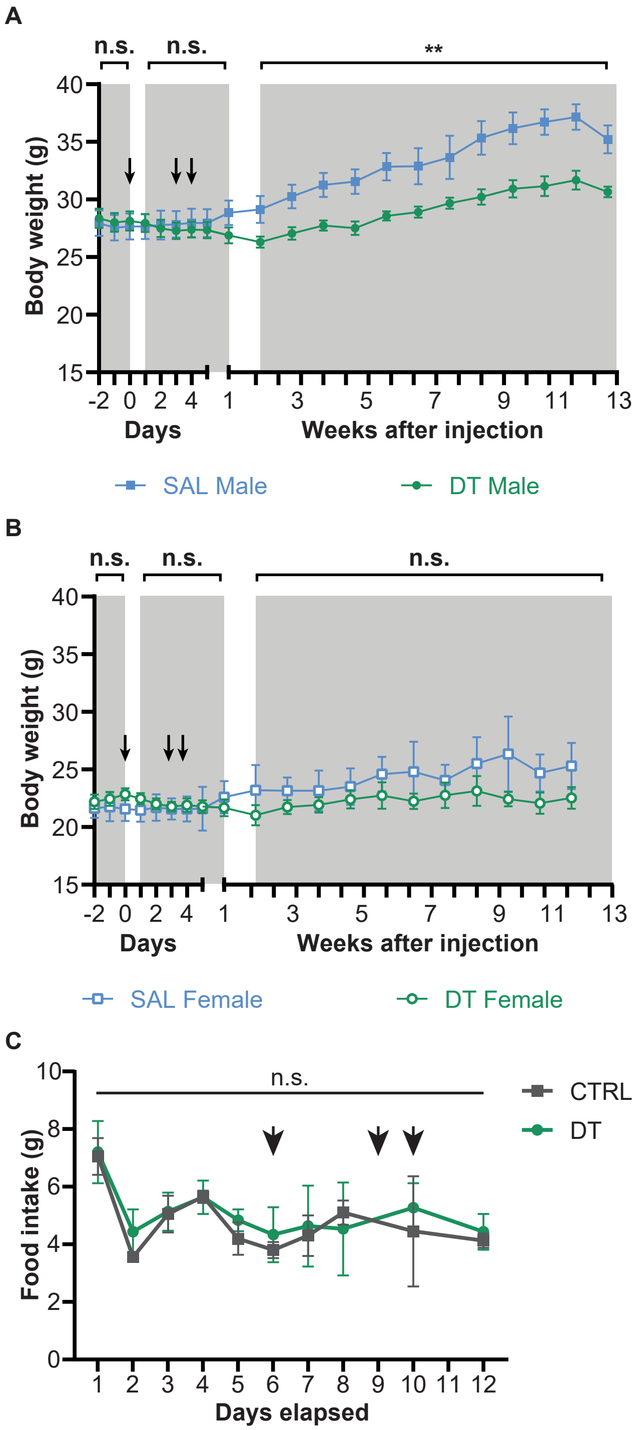


**Figure S2. Body weight and feeding measurements in *Sst*-Cre x lsl-DTR mice, related to Figure 2.** A) Body weight measurements in male SAL-treated (n = 4) and DT-treated (n = 6) *Sst*-Cre x lsl-DTR mice. Black arrows represent IP administration of SAL or DT. B) Body weight measurements in male SAL-treated (n = 3) and DT-treated (n = 3) *Sst*-Cre x lsl-DTR mice. Black arrows represent IP administration of SAL or DT. C) Feeding measurements in DT-treated *Sst*-Cre x lsl-DTR (n = 3) and *Sst*-Cre only (n = 3) mice. Black arrows represent IP administration of DT. Scale bars represent SEM, **p < 0.01.

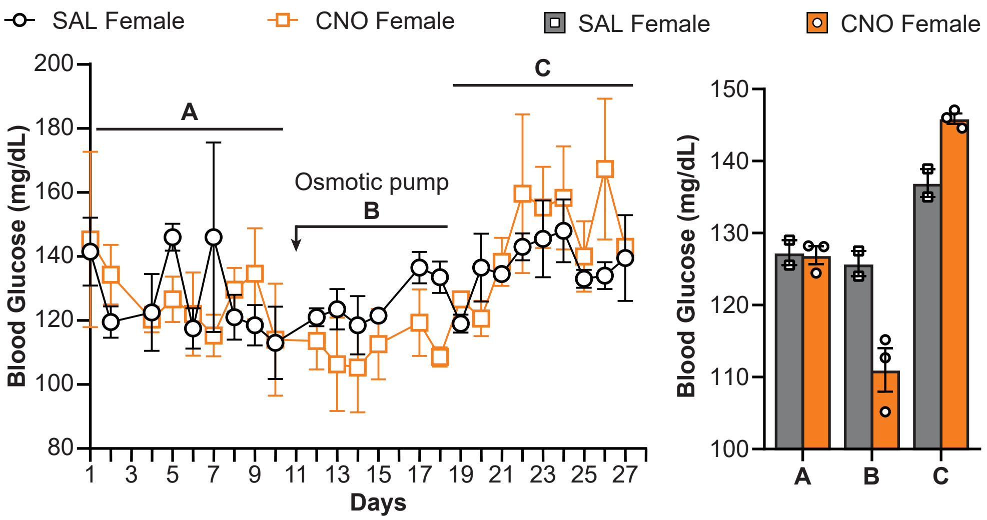


**Figure S3. Body weight and feeding measurements in *Sst*-Cre x lsl-DTR mice, related to Figure 4.** A) Glucose measurements in *Sst*-Cre x lsl-Gi-DREADD females before, during, and after implantation of osmotic pump containing 0.9% saline (n = 2) or 1 mg/mL CNO diluted in 0.9% saline (n = 3). Bar graphs represent average glucose levels of each mouse during the time period specified (A = before implantation, B = during osmotic pump implantation, C = after osmotic pump runs out of CNO). Error bars represent SEM.

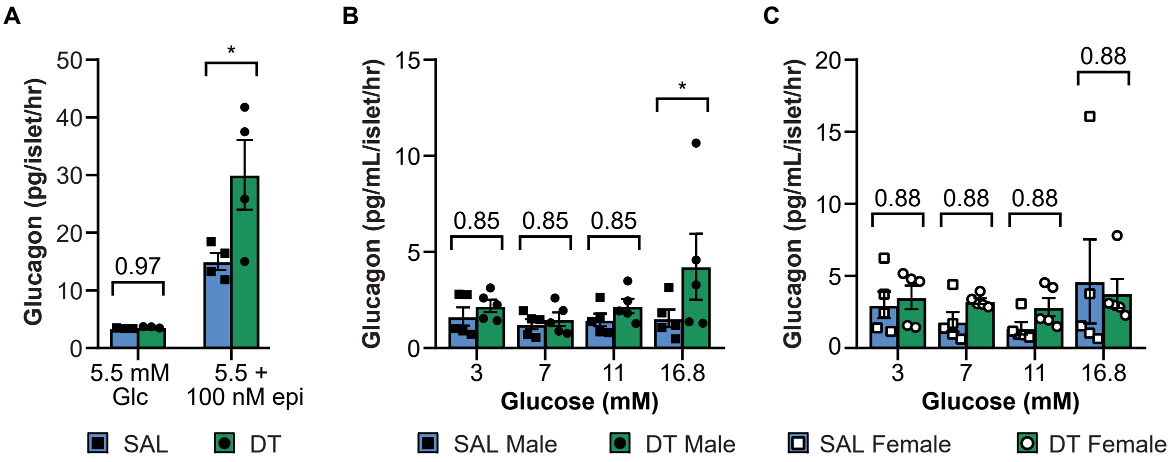


**Figure S4. Body weight and feeding measurements in *Sst*-Cre x lsl-DTR mice, related to Figure 5.** A) Static glucagon secretion assay performed on islets isolated from SAL- or DT-treated *Sst*-Cre x lsl-DTR mice. Islets were stimulated with epinephrine to stimulate glucagon secretion. B) Static glucagon secretion assay performed on the same islets from Figure 3I (male SAL- or DT-treated *Sst*-Cre x lsl-DTR mice). C) Static glucagon secretion performed on the same isletes from Figure 3J (female SAL- or DT-treated *Sst*-Cre x lsl-DTR mice). Error bars represent SEM, *$p < 0.05$.

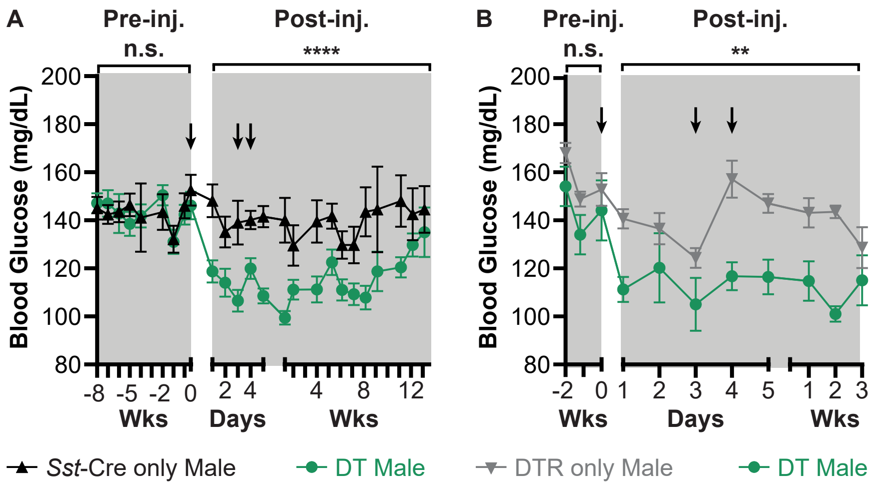


**Figure S5. Glucose measurements in DT-treated *Sst*-Cre only, DTR only, and *Sst*-Cre x lsl-DTR mice, related to Figure 6.** A) Blood glucose measurements of DT-treated male *Sst*-Cre only (n = 5) and *Sst*-Cre x lsl-DTR (n = 10) mice. Black arrows represent IP administration of DT. B) Blood glucose measurements of DT-treated male lsl-DTR only (n = 6) and *Sst*-Cre x lsl-DTR (n = 5) mice. Black arrows represent IP administration of DT. Error bars represent SEM, **$p < 0.01$, ****$p < 0.0001$.
